# *De novo* biosynthesis of alpinetin enhanced by directed evolution of 5-O-methyltransferase

**DOI:** 10.1101/2025.07.06.663367

**Authors:** Bo Peng, Ziwei Wang, Lili Zhang, Matthew R. Groves, Kristina Haslinger

## Abstract

Alpinetin ((2S)-7-Hydroxy-5-methoxyflavan-4-one) is a natural flavonoid found in various medicinal herbs and is frequently used in Chinese patent medicines. It exhibits a wide range of bioactivities, including anti-inflammatory, cardiovascular protective, lung protective, antiviral, hepatoprotective, and antitumor effects. Alpinetin features a 5-methyl group on the A ring, a rare characteristic among methylated flavonoids. The limited abundance of Alpinetin in plant biomass, its laborious extraction and purification from this biomass, and the toxicity and lack of regio- and chemoselectivity in organic synthesis render its biosynthesis in engineered microbes an attractive alternative.

In this study, we aimed to achieve the *de novo* biosynthesis of alpinetin in *Escherichia coli*. In a series of optimization steps, we varied the selection of pathway enzymes, plasmid configurations, medium composition, and fermentation strategies. Lastly, we applied a directed evolution campaign to the 5-O-methyltransferase to successfully enable the *de novo* biosynthesis of alpinetin in *E. coli* for the first time. Moreover, we demonstrated the ability of O-methyltransferase to methylate a broad range of flavonoid substrates, leading to the production of valuable O-methylated flavonoids. Our study represents the first example of alpinetin biosynthesis in a heterologous host and paves the way to produce other valuable O-methylated flavonoids enzymatically.

## 1. Introduction

Pharmacological effects of plant-derived flavonoids have increasingly drawn attention to herbal medicines as alternative treatments for human diseases such as cancer, epilepsy, inflammation, central nervous system (CNS) disorders, diabetes, cardiovascular disorders, and to improve the immune system.^1–7^ Alpinetin (7-Hydroxy-5-methoxyflavan-4-one), a rare flavonoid sourced from *Alpinia galanga* (L.), features an uncommon 5-O-methylation of the A ring. Alpinetin has been found to exhibit anti-inflammatory and antioxidant properties and has been recognized for its potential benefits in treating non-alcoholic fatty liver disease, high-fat diet-aggravated lung carcinogenesis, lipopolysaccharide/D-Galactosamine-induced liver injury, ulcerative colitis, osteoporosis, Zika virus infection, intestinal inflammation, and certain cancers.^8–14^

In Traditional Chinese Medicine, alpinetin is a key component of patented drugs such as Xingqiwenzhong granule, Fufangcaodoukou tincture, Jianweizhitong tablet, and the Baikoutiaozhong pill.^15^ These medications are used to treat digestive disorders, including epigastric pain, belching, nausea, vomiting, and anorexia.^14,16^ Despite the diverse bioactivities of alpinetin, its production primarily relies on extraction from plants, particularly from the seeds and rhizomes of plants from the *Zingiberaceae* family.^17,18^ However, the low extraction efficiency of the multi-step purification process limits the commercial availability of alpinetin. Recently, Sokolova *et al*. reported that an O-methyltransferase (StrAOMT) from the bacterium *Streptomyces avermitilis* can methylate pinocembrin to generate alpinetin in a low yield *in vitro*, offering an interesting starting point for exploring microbial biosynthesis of alpinetin.^19^

O-methylation is a powerful and common tool to improve pharmacological and physicochemical properties of natural products.^20,21^ For example, homoeriodictyol and hesperetin exhibit improved biological activities and pharmacological properties, including metabolic stability, membrane transport capability, and oral bioavailability, compared to their unmethylated counterparts.^22^ Xanthohumol, a flavonoid with methyl and prenyl modifications, displays a range of biological activities, including cancer prevention, mitigation of diabetes, anti-inflammatory, antioxidant, and immunomodulatory effects.^23,24^ The chemical synthesis of alpinetin has not yet been established, and in organic chemistry, methylation reactions typically use methyl iodide, a toxic and unsustainable reagent that is harmful to the environment.^25^ Furthermore, traditional chemical methylation of flavonoids often requires the use of protecting groups for the non-target hydroxyl groups to achieve selectivity, adding to the number of synthesis steps and negatively affecting the atom economy. In contrast, enzymatic methylation with S-adenosyl-L-methionine (SAM) as a methyl donor occurs under mild conditions and potentially with high regio- and chemoselectivity. *In vitro*, SAM analogs can even be applied to facilitate the transfer of a range of natural and non-natural alkyl groups to acceptor substrates.^20,26^

In flavonoid biosynthesis, O-methyltransferases (OMTs) can be classified into two categories: Class I OMTs, which are Mg^2+^-dependent and have a low molecular weight (23-30 kDa), and Class II OMTs, which are high molecular weight (36-43 kDa) and are metal-independent.^27^ In principle, O-methylation can occur at any position; however, 4’- and 7-methylation are the most highly represented^28^ as the 4’-OH group on the B ring is more reactive due to resonance stabilization from the aromatic ring. Meanwhile, the 7-OH group on the A ring is frequently methylated because it is less sterically hindered and more accessible. The 5-hydroxyl group is only rarely modified by methylation or glycosylation because it can form a hydrogen bond with the carbonyl group at carbon 3 and is therefore not very reactive (Figure 1).^29^

**Figure 1.**
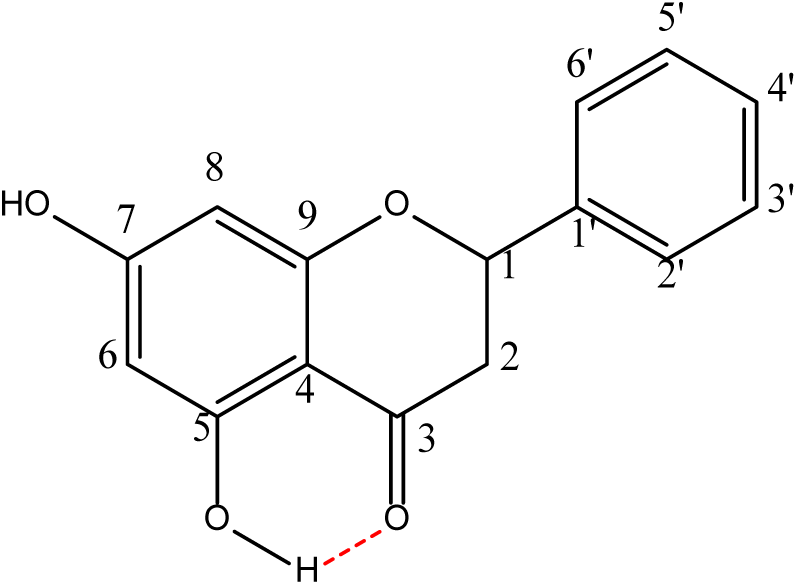
The chemical structure of pinocembrin, the alpinetin precursor. The hydrogen bond between the 5-hydroxy group and the 3-carbonyl group is shown in red.

In this study, we designed a biosynthetic pathway for *de novo* alpinetin production in *Escherichia coli* and successfully enhanced the activity of O-methyltransferase and final titer of alpinetin through directed evolution and fermentation optimization. Further studies using various flavonoids as substrates revealed that StrAOMT and its variants can O-methylate a broad range of flavonoids with specific regioselectivity for certain compounds. Our research represents the first instance of alpinetin biosynthesis in *E. coli* and will pave the way for producing valuable O-methylated flavonoids.

## 2. Materials and methods

### 2.1. Bacterial strains, primers, and plasmids

All bacterial strains and plasmids used in this study are listed in Table S1 and 2. Primers are given in Table S1. *E. coli* DH5alpha (New England Biolabs) was used for routine cloning and plasmid amplification, *E. coli* MG1655 K-12 (DE3) and *E. coli* BL21 (DE3) were used for protein expression and fermentation.

The genes coding for HvCHS, chalcone synthase from *Hordeum vulgare* (GenBank accession number XP_044963194); Os4CL, 4-coumarate-CoA ligase from *Oryza sativa* (GenBank accession number NP_001396278); PhCHS, chalcone synthase from *Petunia hybrida* (GenBank accession number KP284563.1); Pc4CL, 4-coumarate-CoA ligase from *Petroselinum Crispum* (GenBank accession number KX671122.1). StrAOMT, O-methyltransferase from *Streptomyces avermitilis*. MsCHI, chalcone isomerase from *Medicago sativa* (GenBank accession number P28012) are described in our previous studies.^22^ The genes encoding RmXAL (GenBank accession number KR095285), aromatic amino acid ammonia lyase from *Rhodotorula mucilaginosa*; Gm4CL, 4-coumarate-CoA ligase from *Glycine max* (GenBank accession number X69955) and CsCHS, chalcone synthase from *Camellia sinensis* (GenBank accession number: D26593) were codon-optimized and synthesized (sequences provided in the Supporting Information) with the introduction of a restriction site (TWIST Bioscience, San Francisco, CA, USA).

The gene RmXAL was cloned between the BamHI and HindIII restriction sites and the gene MsCHI was cloned between the NdeI and XhoI restriction sites of pACYCDuet-1 (Novagen) yielding pACYCDuet::RmXAL-MsCHI and pACYCDuet::MsCHI, respectively. The gene for CsCHS1 was cloned between the SacI and HindIII restriction sites and the gene Gm4CL was cloned between the NdeI and XhoI restriction sites of pETDuet-1 (Novagen) yielding pETDuet::CsCHS1-Gm4CL, respectively. The gene StrAOMT was cloned between the NcoI and HindIII restriction sites of pRSFDuet-1 and pCDFDuet-1 (Novagen) yielding pRSFDuet::StrAOMT and pCDFDuet::StrAOMT, respectively. The plasmids pETDuet::PhCHS-Pc4CL, pETDuet::HvCHS-Os4CL, and pET21b::StrAOMT have been described previously.^19,22^ All plasmids were isolated with the QIAprep Spin Miniprep kit (QIAGEN) and were verified by Sanger sequencing.

### 2.2. Construction of Recombinant *E. coli* Strains

To create *E. coli* strains s1–s20 capable of producing flavonoids, a series of plasmid combinations were introduced into *E. coli* MG1655 (DE3) and *E. coli* BL21 (DE3) cells using electroporation. The method for preparing electrocompetent *E. coli* was adopted from Green and Sambrook.^30^ After transformation, cells were cultured on selective Luria-Bertani (LB) agar. Depending on the experiment, the culture medium was supplemented with either 100 µg/L ampicillin, 20 µg/L chloramphenicol, 50 µg/L spectinomycin, 50 µg/L kanamycin, or combinations of these antibiotics to provide selective pressure for plasmid maintenance. The cells were incubated overnight at 30 °C for *E. coli* MG1655 (DE3) or at 37 °C for *E. coli* BL21 (DE3). The following day, colonies were transferred to 3 mL of selective LB liquid medium in round-bottom polystyrene tubes and cultivated at 30 °C or 37 °C and 200 rpm overnight. Subsequently, glycerol stocks (containing 50% v/v glycerol) were prepared from these cultures and stored at −70 °C for future use.

### 2.3. Multiple Sequence Alignment and docking study

Protein sequences were aligned with Clustal Omega (1.2.4) with default settings.^31^ The alignment was visualized with ESPript.^31^ Molecular docking was performed using PLANTS1.2 (https://github.com/discoverdata/parallel-PLANTS).^32–34^ Receptor and ligand files were prepared using Open Babel version 3.1.0 and Pymol (Schrödinger, LLC).^35^ The binding site was defined, and re-docking validation was conducted using the crystal structure of the *Myxococcus xanthus* OMT, SafC, in complex with S-adenosylhomocysteins (SAH), Mg^2+^, and L-dopamine (LDP) (PDB ID 5LOG) as a template.^36^ Docking of pinocembrin to StrAOMT (PDB ID 8C9S^19^) was subsequently performed using the same parameterization.

### 2.4. Library construction

The plasmid of pET21b::StrAOMT has been described previously.^19^ Primers used for mutagenesis are listed in Supporting Table S1. Site-saturation mutagenesis was performed using QuikChange with degenerate primers of 32 codon combinations obtained from Eurofins Scientific (Luxembourg). For the PCR, 50 ng of pET21b-StrAOMTserved as the DNA template. The whole plasmid was amplified with Q5 high-fidelity DNA polymerase (New England Biolabs, Frankfurt am Main, Germany) in 50 µL reaction volume. Subsequently, 1 µL Dpn1 was added and incubated at 37 °C overnight. The resulting mixture was transformed into *E. coli* DH5α competent cells and transformants were selected on LB-agar plates, supplemented with 100 µg/mL ampicillin. Plasmids from five randomly selected colonies of each library were isolated and sequenced to ensure library quality. All other colonies were resuspended together in LB medium and plasmid DNA was isolated and used to generate the mutant library in *E. coli* BL21 (DE3).

### 2.5. Screening of StrAOMT variants using cell-free extract

Screening of StrAOMT variants was performed with cell-free extract (CFE) in a 96-deep well format. The library plasmid DNA was transformed into electroporation competent *E. coli* BL21(DE3) and grown on LB agar plates supplemented with 100 µg/mL ampicillin at 37 °C overnight. Bacterial colonies were picked from these agar plates with sterile toothpicks and used to inoculate 125 µL LB medium with 100 µg/mL ampicillin in a 96-well plate. As control, three wells were inoculated with *E. coli* BL21 (DE3) cells harboring pET21b::StrAOMT, pET21b::StrAOMT (K212R), pET21b::StrAOMT (I41L/K212R), or pET21b::StrAOMT (R171W/K212R). The plates were sealed with sterile gas-permeable seals (Breathe-Easy, Diversified Biotech, Boston, MA, USA) and starter cultures were grown at 37 °C, 220 rpm overnight. From this overnight culture, 25 µL were used to inoculate 1.2 mL TB medium supplemented with 100 µg /mL ampicillin in 96-deep well plates that were placed at 37 °C, 220 rpm until an optical density ((λ = 600 nm); OD_600_) of 0.4-0.6. Isopropyl-β-D-1-thiogalactopyranoside (IPTG, final concentration, 1 mM) was subsequently added for induction of protein expression at 30 °C, 220 rpm overnight. The remainder of each overnight culture was used to prepare glycerol stocks stored at - 70 °C for later use. After overnight incubation, the cells in 96-deep well plates were collected by centrifugation (30 min, 3,100 x g), the supernatant was removed and the cell pellets were frozen for 45 min at - 70 °C. The cells were then thawed and incubated for 45 min at 30 °C in 200 µL lysis buffer containing 25 mM Tris-HCl pH 7.5, 1 mg/mL lysozyme and 5 µg/mL Benzonase (Novagen). The insoluble cell fraction was removed by centrifugation (45 min, 3,100 x g) and 90 µL of the CFE was transferred into a 96-deep well plate for activity screening of StrAOMT variants. Enzymatic reactions were performed with 90 µL CFE and 10 µL of a reaction mix including 1 mM pinocembrin, 4 mM S-adenosylmethionine (SAM), 20 mM MgCl_2_ for 6 h at 37 °C. The reactions were quenched with 100 µL methanol and insoluble proteins were removed by centrifugation (45 min, 3,100 x g). The supernatants were transferred into a new 96-well plate for reverse-phase HPLC analysis.

### 2.6. *E. coli* whole-cell biotransformation experiments

For 48 well plate fermentations, the alpinetin-producing strains were streaked in triplicate from glycerol stocks onto selective LB agar plates. After overnight incubation at 30 °C, single colonies were inoculated into starter cultures of 3 mL of selective LB medium in round-bottom polystyrene tubes and grown with shaking at 200 rpm at 30 °C overnight. The next day, the cultures were diluted 1:100 into 3 mL seed cultures of modified selective 4-morpholinepropanesulfonic acid (MOPS) medium for 24 h at 30 °C. These were used to inoculate working cultures of 1 mL at an OD_600_ of 0.05 in a 48 flower-shaped well plate (m2p-labs, Germany) and incubated at 30 °C, 900 rpm in an Eppendorf Thermomixer. Isopropyl-β-d-1-thiogalactopyranoside (IPTG) (final concentration, 1 mM), methionine (final concentration, 0-10 mM), phenylalanine (final concentration, 0.1-3 mM), and cerulenin (final concentration, 20 μg/mL) were added to the cell culture after 0-6 h, and incubation was continued for 36 h. Modified mops medium composition (1x) prepared from sterile stocks: MOPS medium, Trace Mineral Supplement (ATCC MD-TMS, used as 200 x stock), vitamin mix (from 100x stock; final: riboflavin 0.84 mg/L, folic acid 0.084 mg/L, nicotinic acid 12.2 mg/ L, pyridoxine 2.8 mg/L, and pantothenic acid 10.8 mg/L), biotin (from 1000 x stock; final: 0.24 mg/L), thiamine (from 1470 x stock; final: 340 mg/L), 4% (w/v) glucose (from 50% (w/v) stock), MgCl_2_ (from 500x sterile stock in water, final 2 mM).

For fermentation in shake flasks, three single colonies of each respective strain were inoculated into selective LB medium and incubated overnight at 37 °C. The next day, 1 mL of the starter culture was inoculated into 50 mL of fresh, selective TB medium. IPTG (final concentration, 1 mM) was added to the culture broths when the OD_600_ reached 0.4−0.6 for protein expression and cultures were subsequently incubated at 30 °C for 16 h. The cells were collected by centrifugation for 30 min at 3,100 x g (Eppendorf 5920R, Germany) and 4 °C. Two cell pellets were combined and resuspended in 5 mL of fresh modified MOPS medium, which included 1 mM phenylalanine, 0.1 mM methionine, 1 mM IPTG, 4% (w/v) glucose (from 50% (w/v) stock), and 2 mM MgCl_2_ for a further 36 h of fermentation at 30 °C. Samples were taken for quantification of extracellular metabolites after 36 h of fermentation. To determine the concentration of cinnamic acid, pinocembrin, and alpinetin, 1 mL of culture was taken from the flask and mixed with 1 mL of methanol solution (100% methanol with 0.1% (v/v) TFA). The mixture was then centrifuged for 10 minutes at 10,000 x g, and the supernatant was used for HPLC analysis. To analyze the alpinetin content in the fermentation broth, 2 mL samples were taken from the flask and extracted twice with 4 mL of ethyl acetate. The organic phase was collected and evaporated. After evaporation, 1 mL of 50% (v/v) methanol (in water) was added to dissolve the crude extract, which was then filtered using a 0.45 μm PTFE syringe filter to remove insoluble particles before LC-MS analysis. Samples were stored at −20 °C until analysis.

### 2.7. Analysis of target compounds

The authentic standards for pinocembrin, alpinetin, and S-adenosylmethionine (SAM) were purchased from BLDpharm (Shanghai, China). Fatty acid synthase (FAS) inhibitor cerulenin was purchased from Enzo Life Sciences (USA). Cinnamic acid, caffeic acid, phenylalanine, thiamine, kaempferol, luteolin, quercetin, chrysin, naringenin, hesperetin, eriodictyol, homoeriodictyol, phloretin, prunin, naringenin 5-methyl ether, and biotin were purchased from Sigma-Aldrich (USA). Amentoflavone, myricetin, eupatorin were purchased from Extrasynthese (France).

Quantification of cinnamic acid, pinocembrin, and alpinetin from the enzymatic reactions and fermentation broths was achieved by high-performance liquid chromatography (HPLC), with a Shimadzu LC-10AT system equipped with a SPD-20A photodiode array detector (PDA). The samples were analyzed using 20 μL injections and separation over an Agilent Eclipse XDB-C18 (5 μm, 4.6×150 mm) column. Samples from enzymatic reactions were separated by isocratic elution with 45 % mobile phases A (water + 0.1% (v/v) trifluoroacetic acid (TFA)) and 55 % B (acetonitrile + 0.1% (v/v) TFA) at a flow rate of 1 mL/min. To separate metabolites from 48 well plate and flask fermentation, the following gradient was used: 15% B for 3 min, 15-90% B over 6 min; 90% B for 2 min; 90−15% B over 3 min, 15% B for 4 min; flow rate: 1 mL/min. Cinnamic acid, pinocembrin, and alpinetin were identified by comparison to authentic standards. The peak areas were integrated and converted to concentrations based on calibration curves with the authentic standards (Figure S1).

The identity of alpinetin and other O-methylated flavonoid derivatives were assessed by liquid chromatography−mass spectrometry (LC-MS) with a Waters Acquity Arc HPLC-MS system equipped with a 2998 PDA detector and a QDa single-quadrupole mass detector. Samples (2 μL) were injected into and separated over an Xbridge BEH C18 (3.5 μm, 2.1 × 50 mm) column with the following mobile phases: A: water +0.1% (v/v) formic acid (FA); B, acetonitrile +0.1% (v/v) FA. The following gradient was used: 5% B for 2 min, 5−90% B over 3 min, 90% B for 2 min, 90−5% B over 3 min; flow rate: 0.5 mL/min. MS analysis was carried out in positive ionization mode, with the following parameters: probe temperature of 600 °C; capillary voltage of 1.0 kV; cone voltage of 15 V; scan range 100−1250 m/z.

The identity of alpinetin and other O-methylated flavonoid derivatives was further assessed by High-resolution tandem MS analyses, which were performed with a Shimadzu Nexera X2 HPLC system with binary LC20ADXR interfaced to a QExactive Plus Hybrid Quadrupole-Orbitrap Mass Spectrometer (Thermo Scientific). The column, injection volume, flow rate, mobile phases, and LC method were used as described above for HPLC-MS. MS and MS/MS analyses were performed with electrospray ionization in positive mode at a spray voltage of 3.5 kV, a sheath gas pressure of 60 psi, and an auxiliary gas flow of 11 arbitrary units. The ion transfer tube temperature was 300 °C. Spectra were acquired in data-dependent mode with a survey scan at m/z 100−1650 at a resolution of 70,000 followed by MS/MS fragmentation of the top 5 precursor ions at a resolution of 17,500. A normalized collision energy (NCE) of 35 or a mix of 30, 40, and 55 were used for fragmentation, and fragmented precursor ions were dynamically excluded for 10 s.

### 2.8. Protein Expression, Purification, and Activity tests

#### 2.8.1. Expression

*E. coli* BL21(DE3) harboring the plasmids pET21b::StrAOMT, pET21b::StrAOMT (K212R), pET21b::StrAOMT (I41L/K212R), pET21b::StrAOMT (R171W/K212R), pET21b::StrAOMT (I41L/R171W/K212R) were inoculated in 3 mL LB (containing 100 μg/mL ampicillin) for overnight cultivation at 37 °C and 200 rpm. The next day, 1 mL of overnight cell culture was used to inoculate 1 L of selective TB medium at 37 °C and 200 rpm in a 5 L glass Erlenmeyer flask. When the OD_600_ reached 0.4-0.6, 1 M IPTG (final concentration, 1mM) was added for induction of protein expression. The TB medium composition (1 L) consisted of 20 g tryptone, 24 g yeast extract, 4 g glycerol, 2.3 g potassium dihydrogen phosphate, 12.5 g dipotassium phosphate, and 100 μg/mL ampicillin.

#### 2.8.2. Purification

All steps were performed with cooled buffers at 4 °C. Cells were harvested by centrifugation at 3,100 x g for 20 min. The cell pellet was resuspended in 20 mL of lysis buffer (50 mM Tris−HCl and 200 mM NaCl, pH 7.5). The cells were disrupted by sonication for 4 × 40 s (with a 5 min rest interval between each cycle) at 60 W output. The unbroken cells and debris were removed by centrifugation (10,000 x g for 1 h). The supernatant was filtered through a syringe filter (pore diameter, 0.45 μm) and incubated with 3 mL of Ni^2+^-NTA sepharose resin, which had previously been equilibrated with lysis buffer, in a small column at 4 °C for 18 h with agitation. The unbound proteins were eluted from the column using gravity flow. The column was first washed with lysis buffer (15 mL) and then with buffer A (30 mL, 50 mM Tris−HCl, 200 mM NaCl, and 30 mM imidazole, pH 7.5). Retained proteins were eluted with buffer B (5 mL, 50 mM Tris−HCl, 300 mM NaCl, and 500 mM imidazole, pH 7.5). Fractions were analyzed by separation using sodium dodecyl sulfate polyacrylamide gel electrophoresis (SDS-PAGE)(4−12% polyacrylamide) and staining with the InstantBlue Coomassie Protein stain (Abcam, UK). Fractions containing O-methyltransferase were pooled and loaded onto a HiLoad 16/600 Superdex 200 pg column, which had previously been equilibrated with buffer C (180 mL, 10 mM HEPES, 200 mM NaCl buffer, 5% glycerol, and 10 mM DTT, pH 7). Elution was performed in buffer C at 1 mL/min for 1.2 column volumes. Fractions were collected and analyzed by SDS-PAGE. The purified enzymes were concentrated with centrifugal devices with Omega membrane 30K (Pall, USA) and stored at −70 °C until further use.

### 2.9. Activity tests

The *in vitro* OMT reactions were conducted in a buffer containing 25 mM Tris-HCl (pH 7.5), 2 mM MgCl₂, 1 mM SAM, 0.5 mM substrate, and 5 μM enzyme in a total reaction volume of 100 μL. Reactions were incubated at 37 °C for 6 hours. For kinetic characterization of StrAOMT wild-type and the StrAOMT I41L/R171W/K212R variant, reactions were performed under similar conditions with substrate concentrations ranging from 6.25 to 625 μM and 50 μM enzyme in a total volume of 100 μL. Incubations were carried out at 37 °C for 20, 40, and 60 minutes. Substrate scope screening was conducted in a buffer containing 25 mM Tris-HCl (pH 7.5), 2 mM MgCl₂, 1 mM SAM, 500 μM substrate, and 50 μM enzyme in a final volume of 100 μL, with incubation at 37 °C for 12 hours. Reactions were initiated by the addition of enzyme (or water for the “no OMT” control), incubated at 37 °C, then quenched with 100 μL 100% methanol containing 0.1% (v/v) TFA. Samples were centrifuged and stored at -20 °C until analysis.

## 3. Result

### 3.1 Biosynthesis of alpinetin

Several studies have documented the production of pinocembrin, the precursor of alpinetin, in *E. coli* and *Saccharomyces cerevisiae*.^37–41^ It can be obtained from phenylalanine via several enzymatic steps catalyzed by aromatic amino acid ammonia lyase (XAL), 4-courmate: CoA ligase (4CL), chalcone synthase (CHS), and chalcone isomerase (CHI), respectively. Based on the results of previous studies on cinnamic acid production in *E. coli*, ^42^ we chose the aromatic amino acid ammonia lyase from *Rhodotorula mucilaginosa* (RmXAL) for cinnamic acid production. To produce pinocembrin chalcone, we tested different combinations of CHS and 4CL enzymes, namely PhCHS and Pc4CL, known for their high efficiency of naringenin production^43^; HvCHS and Os4CL, previously utilized for synthesis of methylated flavonoids^22^; and CsCHS1 and Gm4CL, previously recognized for their capacity to generate significant amounts of pinocembrin.^39^ Lastly, we used chalcone isomerase from *Medicago sativa* for cyclization of pinocembrin chalcone and the recently characterized O-methyltransferase from *Streptomyces avermitilis*, StrAOMT (Figure. 2).^19^

**Figure 2.**
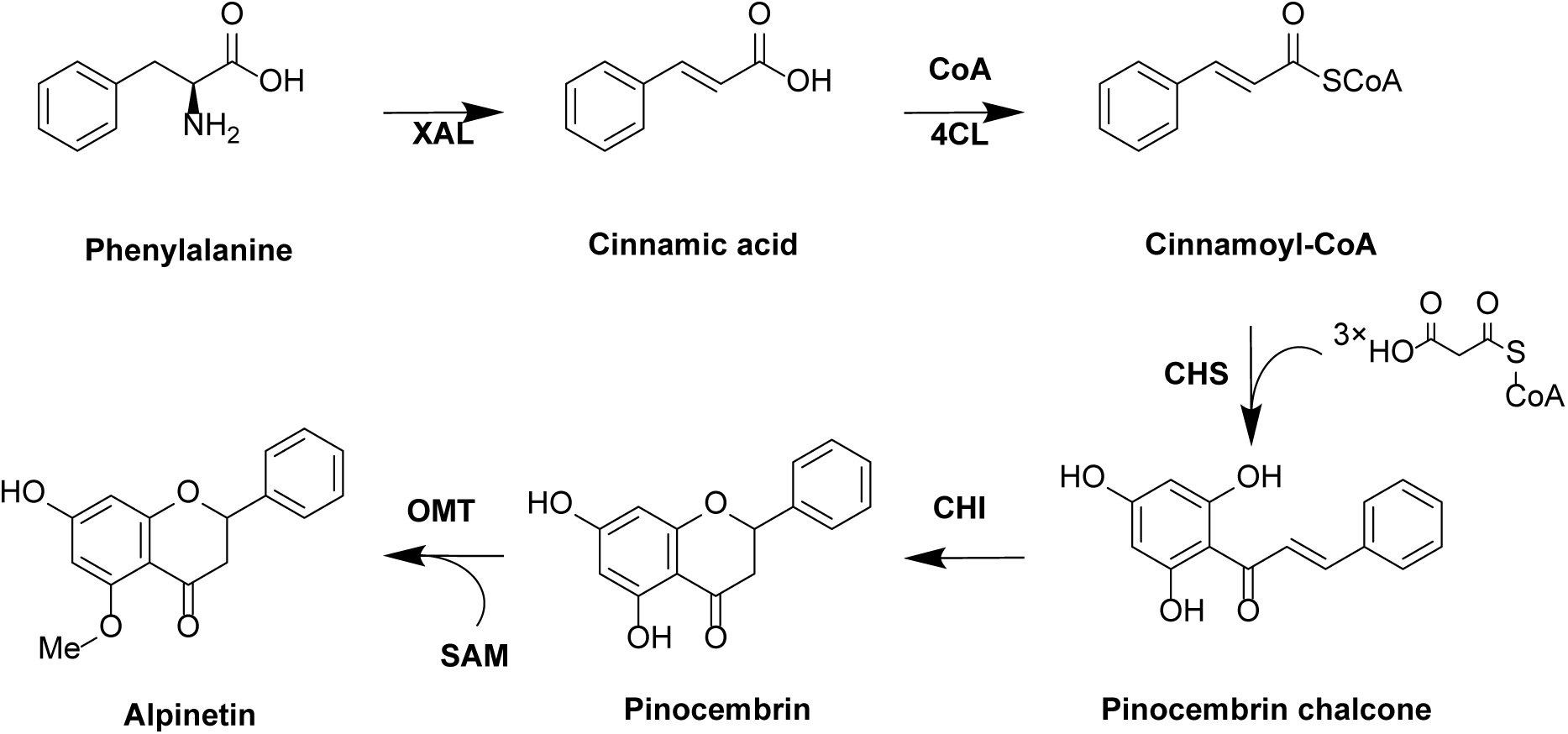
The proposed biosynthesis pathway for alpinetin. XAL, aromatic amino acid ammonia lyase, 4CL, 4-coumarate: CoA ligase; CHS, chalcone synthase; CHI, chalcone isomerase, OMT, O-methyltransferase.

Strains s1-s3 were constructed by co-transformation of the resulting plasmids into *E. coli* MG1655 (DE3) (see Table 2). Initial fermentations were carried out in 1 mL modified MOPS minimal medium, supplemented with 3 mM phenylalanine. The results indicated that the combination of PhCHS and Pc4CL (s1) produced the highest titer of pinocembrin (approximately 56 μM (Figure 3A)). In contrast, combining HvCHS and Os4CL (s2) resulted in a high titer of cinnamic acid but low titer of pinocembrin (0.5 μM). Additionally, the combination of CsCHS1 and Gm4CL (s3) produced around 27 μM of pinocembrin, but no alpinetin was observed in any of the fermentations. Since s2 produced less pinocembrin compared to the other strains, the combination of HvCHS and Os4CL was excluded from further experiments.

**Table 1.**
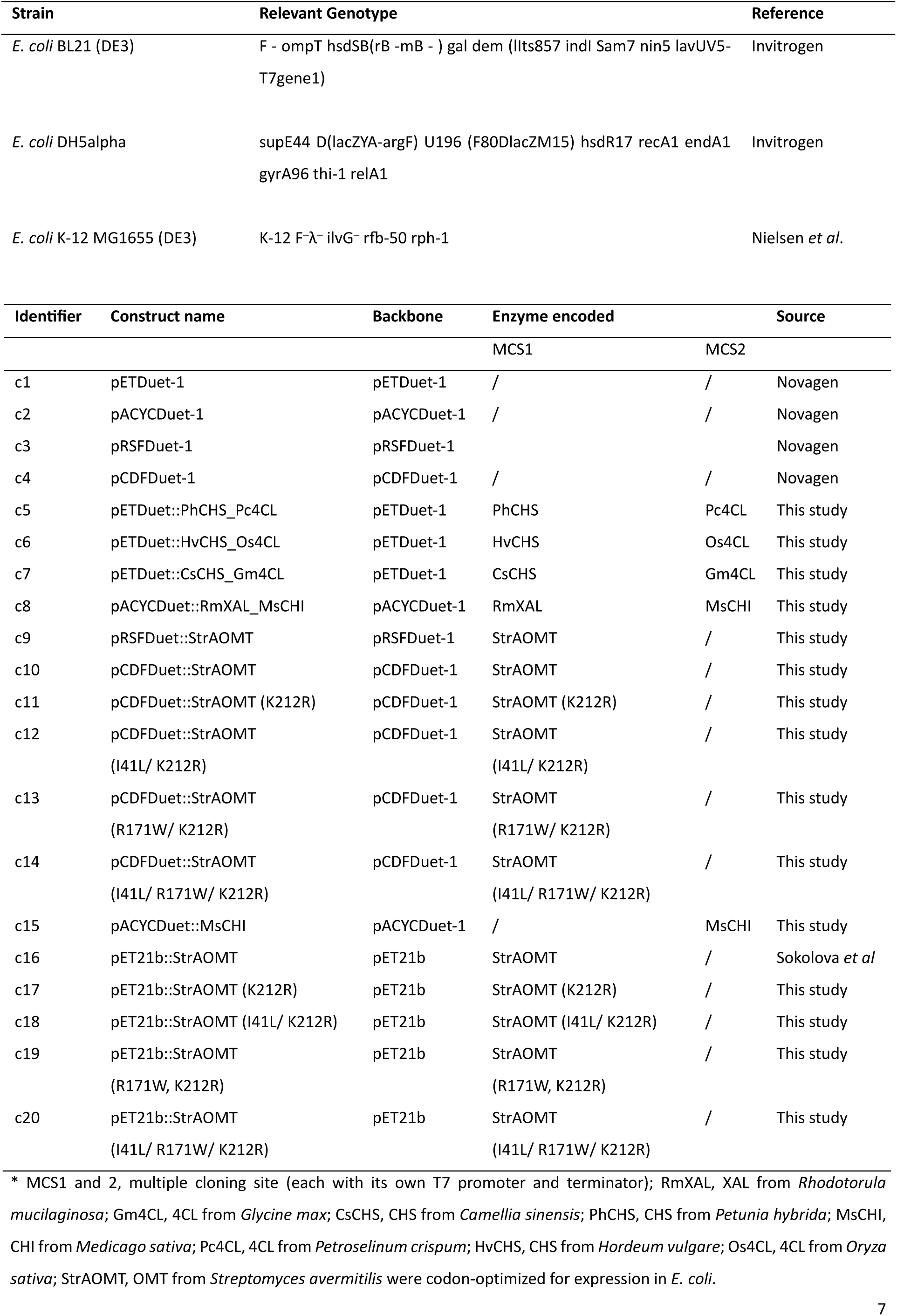
List of bacterial strains and plasmids used in this study.

**Table 2.** List of alpinetin-producing strains used in fermentation experiments.

| Identifier | Construct name | enzymes expressed | *E. coli* strain |
| --- | --- | --- | --- |
| s1 | c5, c8, c9 | RmXAL, Pc4CL, PhCHS, StrAOMT, MsCHI | MG1655 (DE3) |
| s2 | c6, c8, c9 | RmXAL, Os4CL, HvCHS, StrAOMT, MsCHI | MG1655 (DE3) |
| s3 | c7, c8, c9 | RmXAL, Gm4CL, CsCHS1, StrAOMT, MsCHI | MG1655 (DE3) |
| s4 | c5, c8, c9 | RmXAL, Pc4CL, PhCHS, StrAOMT, MsCHI | BL21 (DE3) |
| s5 | c7, c8, c9 | RmXAL, Gm4CL, CsCHS1, StrAOMT, MsCHI | BL21 (DE3) |
| s6 | c5, c8, c10 | RmXAL, Pc4CL, PhCHS, StrAOMT, MsCHI | BL21 (DE3) |
| s7 | c7, c8, c10 | RmXAL, Gm4CL, CsCHS1, StrAOMT, MsCHI | BL21 (DE3) |
| s8 | c1, c2, c10 | StrAOMT | BL21 (DE3) |
| s9 | c4, c5, c8 | RmXAL, Pc4CL, PhCHS, MsCHI | BL21 (DE3) |
| s10 | c4, c7, c8 | RmXAL, Gm4CL, CsCHS1, MsCHI | BL21 (DE3) |
| s11 | c7, c8, c11 | RmXAL, Gm4CL, CsCHS1, StrAOMT (K212R),<br>MsCHI | BL21 (DE3) |
| s12 | c7, c8, c12 | RmXAL, Gm4CL, CsCHS1, StrAOMT (I41L/K212R),<br>MsCHI | BL21 (DE3) |
| s13 | c7, c8, c13 | RmXAL, Gm4CL, CsCHS1, StrAOMT (R171W/K212R),<br>MsCHI | BL21 (DE3) |
| s14 | c7, c8, c14 | RmXAL, Gm4CL, CsCHS1, StrAOMT (I41L/R171W/K212R),<br>MsCHI | BL21 (DE3) |
| s15 | c7, c14, c15 | Gm4CL, CsCHS1, StrAOMT (I41L/R171W/K212R),<br>MsCHI | BL21 (DE3) |
| s16 | c16 | StrAOMT | BL21 (DE3) |
| s17 | c17 | StrAOMT (K212R) | BL21 (DE3) |
| s18 | c18 | StrAOMT (I41L/ K212R) | BL21 (DE3) |
| s19 | c19 | StrAOMT (R171W, K212R) | BL21 (DE3) |
| s20 | c20 | StrAOMT (I41L/ R171W/ K212R) | BL21 (DE3) |

**Figure 3.**
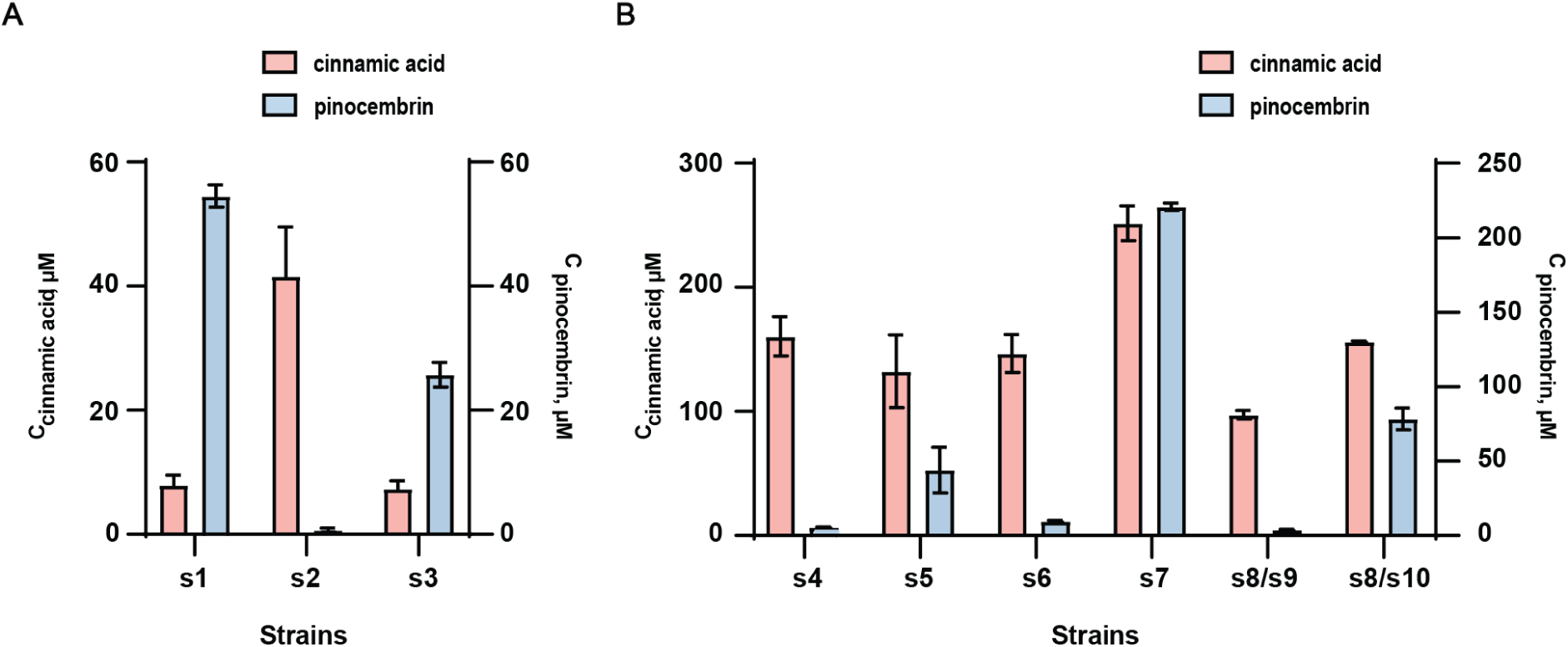
A) Fermentation of *E. coli* MG1655 (DE3) harboring the recombinant flavonoid biosynthesis pathways. Titers of cinnamic acid (red, left axis) and pinocembrin (blue, right axis) were determined 48 h after induction of enzyme expression. B) Biotransformation in resting *E. coli* BL21 (DE3) harboring the recombinant flavonoid biosynthesis pathway. s7 was seen to produce trace amounts of alpinetin, as detected by LC-MS. Bars represent mean +/-SD, n=3.

Subsequently, we conducted a series of experiments for s1 to optimize (co-)substrate (methionine, phenylalanine) feeding time and concentration to enhance pinocembrin production and facilitate the generation of alpinetin. Our findings revealed that a lower concentration of phenylalanine (0.1 mM), the absence of methionine, and a substrate feeding time of 3 hours enhanced the titer of pinocembrin, achieving a highest titer of approximately 92.6 μM (Figure S2). However, alpinetin was still not detected in the final fermentation broth.

Initially, we attributed the lack of alpinetin production to low availability of endogenous SAM *in vivo*, or low concentration of the OMT enzyme. Fermentations were initially performed in MOPS minimal media, which imposes a metabolic burden on *E. coli* and may lead to lower protein expression levels and lower ATP generation. ATP is crucial for SAM biosynthesis as it provides the adenosyl group and serves as an energy donor.^44^ Therefore, we decided to perform the following biotransformations in resting *E. coli* BL21 (DE3) cells to uncouple cell growth and protein expression performed in rich media from the flavonoid production phase performed in modified MOPS media. We co-transformed the plasmids into *E. coli* BL21 (DE3) to generate s4 and s5 and subjected them to the two-step protocol. Although we still did not detect alpinetin in the broth, our experiments revealed that the combination of CsCHS and Gm4CL (s5, 46.5 μM) produced more pinocembrin than the combination of PhCHS and Pc4CL (s4, 6.5 μM) in this strain under these experimental conditions (Figure 3B).

In our initial plasmid configuration StrAOMT was expressed from the high copy-number plasmid pRSFDuet-1, which may impose a metabolic burden according to a previous study.^45^ Therefore, we changed the expression vector of StrAOMT to a lower copy-number plasmid (pCDFduet) to create s6 and s7, which increased pinocembrin production to 14.75 μM (s6) and 205 μM (s7) (Figure 3B). In s7, we found trace amounts of alpinetin detected by low resolution mass spectrometry (around 0.25 μM) and further confirmed by high-resolution tandem MS analyses (Figure S3).

To further alleviate the metabolic burden, we then tested a strategy with two strains. We generated strain s8, expressing only StrAOMT at the highest possible level, and strains s9 and s10, which carried the genes encoding for RmXAL, MsCHI, and either Pc4CL/PhCHS or Gm4CL/CsCHI, respectively. We grew the strains separately for 16 h in 50 mL TB medium after inducing protein expression, harvested them, combined the cell pellets and re-suspended them in 5 mL selective MOPS medium supplemented with 1 mM phenylalanine, 0.1 mM methionine, 1 mM IPTG, 4% (w/v) glucose (from 50% (w/v) stock), and 2 mM MgCl_2_. Despite these efforts, the final titers of pinocembrin were lower than in s6 and s7 and no alpinetin was detected (Figure 3B).

To exclude the possibility that OMT was entirely dysfunctional in our engineered strains, we then tested the ability of s7 and s8 to methylate caffeic acid, a previously reported substrate of StrAOMT with the same two-step biotransformation protocol.^19,21^ We used 1 mM caffeic acid as a substrate and obtained 0.12 mM ferulic acid and 0.036 mM isoferulic acid in the culture broth of s7, and 0.22 mM ferulic acid and 0.04 mM isoferulic acid of s8 (Figure S4). The molar yield of products from s8 is comparable to results obtained in our previous study.^21^ This indicates that StrAOMT is expressed and active in these strains with sufficient SAM supply. Thus, we conclude that StrAOMT is likely the catalyst generating the trace amounts of alpinetin in the fermentation of s7, however, its low affinity and/or catalytic efficiency for this substrate limits product titers.

### 3.2 Site-saturation mutagenesis of 5-O-methyltransferase

After conducting a series of optimizations in the choice of pathway enzymes, plasmid configurations, medium composition, and fermentation strategies, alpinetin production was still very low. We attribute this issue to the low activity of the O-methyltransferase. Previous studies on class I O-methyltransferases have indicated that they carry a conserved catalytic triad of lysine-asparagine-aspartate (K-N-D) and that the adjacent hydroxyl groups of the substrate coordinates the metal ion during catalysis.^19,46,47^

Based on the published crystal structures of StrAOMT in complex with SAH (PDB ID 8C9S)^19^ and substrate-bound SafC (PDB ID 5LOG)^36^, we performed docking experiments using pinocembrin as the ligand.^34^ Ten poses were generated with scores ranging from 252.677 to 260.505. A lower score indicates higher stability of the ligand bound to the protein. After manually inspecting all docking poses, we first identified those that exhibited structural analogy to the experimental SafC structure and satisfied steric complementarity, and then selected from these the pose with the lowest binding score. (Figure 4A). In this pose, hydrogen bonds were observed between pinocembrin and Asp141, Asn168, Lys212, and Mg^2+^. The distance between the side chain amine of Lys144 and the 5-hydroxy group of pinocembrin is 3.5 Å. The 5-hydroxyl group and the 3-carbonyl group of pinocembrin are likely to form a bidentate interaction with the Mg²⁺ ion, stabilizing the substrate within the active site. Under the catalytic action of Lys144, the 5-hydroxyl group undergoes deprotonation, generating a phenolate anion. This deprotonated pinocembrin anion then acts as a nucleophile, attacking the methyl group of SAM in an SN_2_ mechanism, resulting in direct methyl transfer and the formation of the methylated product.^48^ The final model of the substrate-bound StrAOMT was then exported for further analysis. Upon examining residues within 5 Å of pinocembrin, we identified 11 residues: H40, I41, D141, A142, D143, K144, D167, N168, R171, K212, and D215. Notably, D141, D167, and N168 form the metal-binding site in StrAOMT and K144, N168, and D215 the catalytic triad, which is strictly conserved in this OMT family. Next, we performed a protein sequence alignment of SafC (Uniprot accession: Q50859), StrAOMT (Uniprot accession: Q82B68), and NiaKOMT (Uniprot accession: G8T6H8) (Figure 5). In addition to the metal binding and the catalytic residues, positions A142 and D143 were highly conserved among OMTs and were therefore excluded from further mutagenesis. Consequently, the positions H40, I41, R171, and K212 were selected for further analysis and we performed site-saturation mutagenesis for these four positions individually.

**Figure 4.**
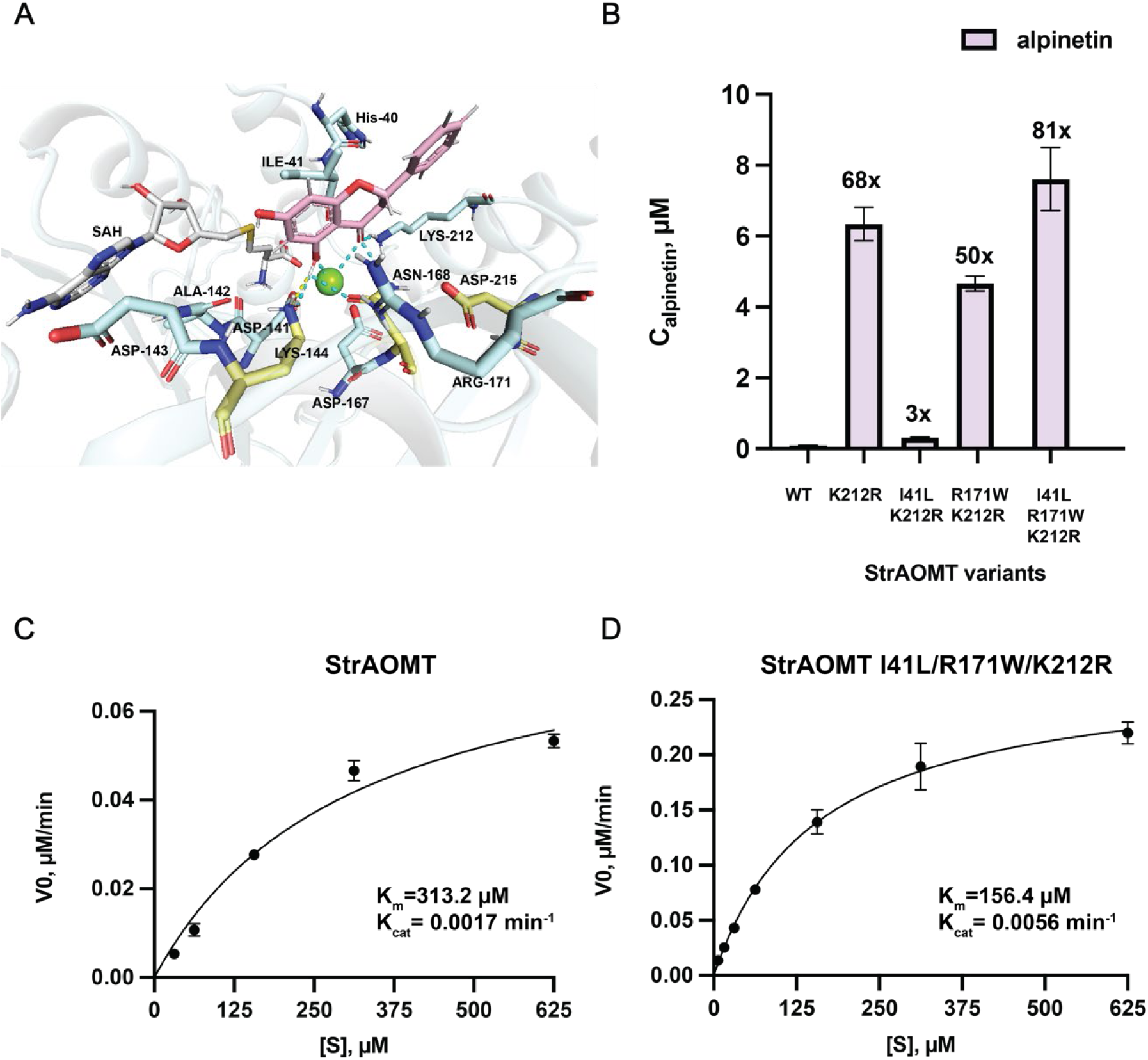
A) Docking model of StrAOMT (PDB ID 8C9S) with pinocembrin (active site residues surrounding the bound SAH and the docked pinocembrin shown as stick representation with atom coloring: blue - nitrogen, red – oxygen, yellow – sulfur, grey - SAH carbon, light blue – StrAOMT carbon, pink – pinocembrin carbon). B) Concentration of alpinetin produced in *in vitro* reactions with purified StrAOMT variants (reaction conditions: 5 μM enzyme, 500 μM pinocembrin, 2 mM SAM, 2 mM Mg^2+^ in Tri-HCL buffer (pH 7.5) at 37 °C for 6 h). C and D) Kinetic characterization of StrAOMT wild type and StrAOMT I41L/R171W/K212R for alpinetin formation. Apparent initial velocities were plotted against varying concentrations of pinocembrin (6.25, 15.625, 31, 62.5, 156, 312.5, and 625 μM), 50 μM purified StrAOMT variants, 2 mM SAM, 2 mM Mg^2+^ in Tri-HCL buffer (pH 7.5) at 37 °C. Data points represent mean ±SD, n = 3. Time-course data used to determine apparent initial velocities are shown in Figure S5.

**Figure 5.**
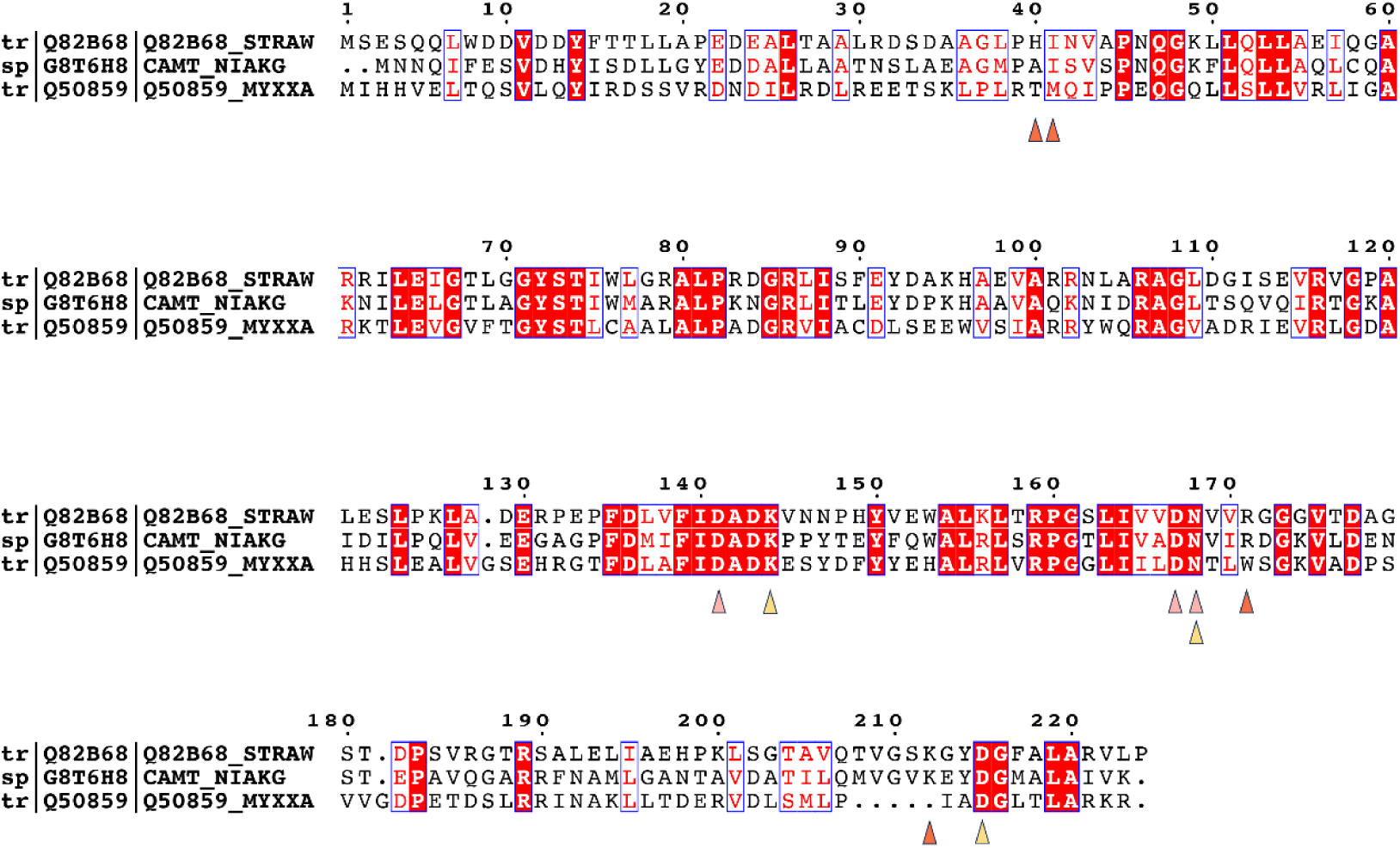
Protein sequence alignment of OMTs (1. O-methyltransferase from *Streptomyces avermitilis*, 2. O-methyltransferase from *Niastella koreensis*, and 3. O-methyltransferase from *Myxococcus xanthus*). Pink triangles indicate the metal-binding sites, yellow triangles indicate the catalytic triad, and orange triangles indicate the residues selected for directed evolution.

The resulting mutant libraries were transformed into *E. coli* BL21 (DE3) and screened by evaluating ∼100 transformants of each library. The screening led to an improved variant of StrAOMT K212R, which exhibited a 4.5-fold increase in alpinetin concentration compared to the wild-type enzyme. To further enhance the overall performance of StrAOMT, we employed an iterative saturation mutagenesis (ISM) strategy, using the best hits from the single-site libraries as templates and randomizing the other respective positions. Consequently, the libraries H40NNK/K212R, I41NNK/K212R, and R171W/K212NNK were constructed. Screening these libraries revealed that the best variant from each variant library were the combinations I41L/K212R, and R171W/K212R, resulting in a 1.6-fold and 2-fold increase in alpinetin concentration after 6 hours compared to the StrAOMT K212R, respectively. However no improved variant was found in the H40NNK/K212R library. We then used StrAOMT I41L/K212R and R171W/K212R as templates to generate the triple mutant libraries StrAOMT I41L /R171NNK/K212R/, and I41NNK/R171W/K212R. Screening these mutants showed that the triple mutant StrAOMT I41L/R171W/K212R achieved a 1.8-fold increase in alpinetin concentration compared to the double mutant variants. We then purified various StrAOMT variants, including the wild-type, K212R, I41L/K212R, R171W/K212R, and I41L/R171W/K212R, for *in vitro* reactions. The results revealed significant increases in alpinetin production compared to the wild type: StrAOMT K212R, R171W/K212R, and I41L/R171W/K212R displayed a 68-fold, 50-fold, and 81-fold increase, respectively. StrAOMT I41L/K212R showed a more modest 3-fold increase (Figure 4B). The results indicate that the K212R mutation has the strongest effect on the interaction with the substrate and/or the catalytic activity of the enzyme and that the I41L mutation is only beneficial in combination with the R171W mutation.

Intrigued by the observation that the triple mutant exhibited 81-fold increase in alpinetin titer, we performed steady-state kinetic assays for StrAOMT wild-type and StrAOMT I41L/R171W/K212R using SAM at a fixed concentration (2 mM) and pinocembrin at variable concentrations. Plotting the apparent initial reaction velocities at different pinocembrin concentrations allowed us to fit the data with the Michaelis−Menten equation (Figures 4C, D, and S5). The v_max_ and k_cat_ of the triple mutant are higher than those of the wild type, showing increases of 3.3-fold and 3.5-fold, respectively. This suggests that StrAOMT I41L/R171W/K212R has a higher turnover number and enhanced enzymatic efficiency. Additionally, the K_M_ of StrAOMT wild type is twice that of the StrAOMT triple mutant, indicating that the StrAOMT I41L/R171W/K212R variant has a higher affinity for pinocembrin compared to the wild type. Still, the affinity of the triple mutant for pinocembrin is rather low, which suggests that OMT may not work optimally *in vivo* because pinocembrin concentrations are too low.

### 3.3 Production of alpinetin in resting cells expressing the enhanced O-methyltransferase variants

After identifying several mutants that showed improved affinity and catalytic activity towards pinocembrin compared to the wild type, we assessed their performances in the context of the flavonoid biosynthesis pathway. We generated the strains s11-s14 harboring these mutants, and carried out biotransformations in resting cells alongside s7 to assess whether the improved variants of StrAOMT enhance alpinetin production. The results showed that strain s7 produced approximately 0.13 μM alpinetin, while the highest alpinetin titer was achieved with strain s14 harboring the triple mutant of StrAOMT (0.96 μM), representing about a 7-fold increase. Still, the majority of the pinocembrin produced in this strain remains unmethylated, which is likely related to the moderate affinity of the OMT enzyme for pinocembrin. However, the poor *in vivo* performance cannot be fully explained by this effect.

To further rationalize our findings, we collected and analyzed extracts from the supernatant and the cell pellet separately and observed that pinocembrin accumulated in the supernatant. We hypothesized that the intracellular concentration of pinocembrin at any given moment is too low for StrAOMT to catalyze the methylation efficiently. As a result, we decided to vary the media composition of the biotransformation step to assess whether supplementation of phenylalanine, glucose and extracellular SAM were beneficial to alpinetin production in the best-performing strain s14 (Figure 6B). The results indicated that phenylalanine supplementation did not impact final alpinetin production, but rather reduced the amount of accumulated pinocembrin (Figure 6). Glucose addition reduced the final titers of both alpinetin and pinocembrin. SAM supplementation increased the final alpinetin titer by 2.9-fold, from 1.71 μM to 4.98 μM. Interestingly, we also detected naringenin (around 5 μM) and 5-methyl-naringenin, a 5-O-methylated flavonoid commonly co-occurring with alpinetin in *Alpinia* genus plants,^49^ at concentrations below 1 μM. The production of 5-methyl-naringenin could be due to the promiscuity of XAL. In Indeed, a previous study showed that XAL from *Rhodotorula mucilaginosa* is capable of producing high concentrations of cinnamic acid from phenylalanine and low concentrations of *p*-coumaric acid from tyrosine.^42^ *P*-coumaric acid can then enter the flavonoid biosynthetic pathway, leading to the production of naringenin, which can subsequently be methylated to form 5-methyl-naringenin. To confirm this hypothesis, we constructed the plasmid c15 lacking the gene encoding XAL and co-transformed it with c7, and c14 to generate strain s15. When fed with 0.1 mM cinnamic acid as substrate, the resting strain under optimal media conditions produced alpinetin at a concentration of 1.5 μM with no naringenin and 5-methyl-naringenin detected (SI Figure S6).

**Figure 6.**
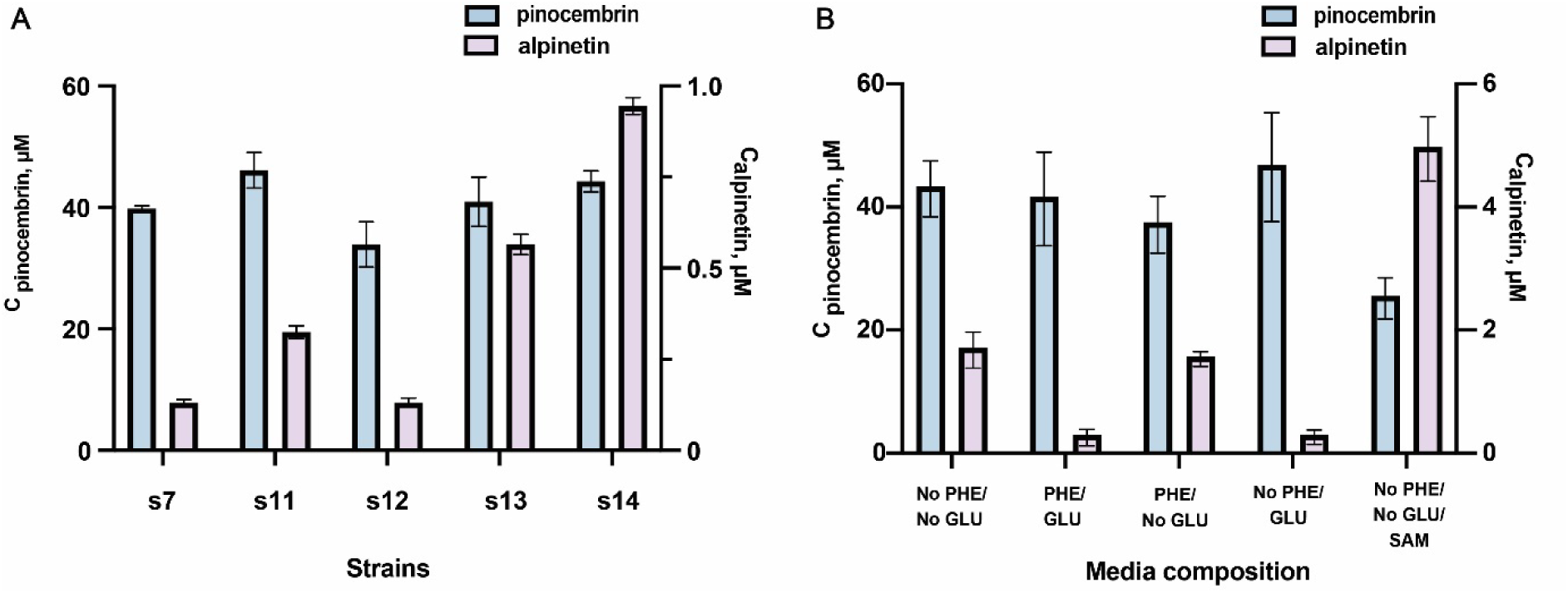
A) Biotransformations of fed phenylalanine to alpinetin in resting *E. coli* BL21 (DE3) harboring the recombinant flavonoid biosynthesis pathways and StrAOMT variants. B) Optimization of the media composition during the resting cell biotransformations with s14. Bars represent mean +/-SD, n=3. Titers of pinocembrin(blue) and alpinetin (ashy purple) were determined after 36 h of incubation.

### 3.4 Substrate scope of StrAOMT variants

Intrigued by the result that the triple mutant variant also produced 5-methyl-naringenin in whole cells, we explored how the substrate scope may have changed compared to our study from 2023.^19^ Thus, we performed in *vitro* experiments with the top two variants (StrAOMT K212R, and StrAOMT I41L/R171W/K212R) and StrAOMT wildtype on other flavonoids: naringenin (**1**), eriodictyol (**2**), homoeriodictyol (**3**), hesperetin (**4**), prunin (a flavanone glycoside, **5**), chrysin (**6**), eupatorin (a trimethoxyflavone, **7**), kaempferol (**8**), luteolin (**9**), and amentoflavone (**10**) (Figure 7). Methylation products were identified based on characteristic shifts in retention times (RT) and an increase in *m/z* values by +14 (single methylation) or +28 (double methylation), relative to the corresponding substrates (Table 3, Figures S6 and S7). Both low-resolution LC-MS and high-resolution tandem MS were used, with data compared against natural product databases and literature reports on O-methylated flavonoids.^50^ In the absence of OMTs, only the unmodified substrate was detected. Using naringenin (**1**, *m/z* 273.0756, [M + H]^+^, RT = 5.62 min) as a substrate, StrAOMT variants produced a single monomethylated product (**1a**) with a retention time of 5.22 min, confirmed as naringenin 5-methyl ether based on a commercial reference standard. For eriodictyol (**2**, *m/z* 289.0706, [M + H]^+^, tR = 6.63 min), two monomethylated products were detected at 7.52 min and 7.67 min, identified as homoeriodictyol (**3**) and hesperetin (**4**), respectively. When homoeriodictyol (**3**, *m/z* 303.0863, [M + H]^+^, RT = 7.50 min) was used as a substrate, two monomethylated products (**3a** and **3b**) were detected at 6.20 min and 8.42 min. MS2 fragmentation analysis suggested that **3a** is methylated on ring A, and its earlier retention time compared to **3** indicates that it could be homoeriodictyol 5-methyl ether. The later-eluting product **3b** exhibited an MS2 fragment pattern consistent with methylation at the 4’-hydroxy group on ring B, identifying it as homoeriodictyol 4’-methyl ether (**3b**), commercially known as hesperetin 3’-methyl ether. For hesperetin (**4**, *m/z* 303.0863, [M + H]^+^, RT = 7.65 min), two monomethylated products were detected at 6.47 min and 8.42 min (**4a** and **4b**, respectively). MS2 analysis of **4a** suggests methylation on ring A, with its earlier retention time suggesting that it is hesperetin 5-methyl ether. The later-eluting product **4b** has the identical RT and M2 fragmentation pattern as **3b**, again supporting its identification as hesperetin 3’-methyl ether. When prunin (**5**, *m/z* 435.1288, [M + H]^+^, RT = 5.17 min) was used as a substrate, two monomethylated products (**5a** and **5b**) were detected at 4.77 min and 5.59 min. MS2 analysis indicated that **5a** is methylated on ring A at the 5-hydroxy position, identifying it as prunin 5-methyl ether. In contrast, **5b** exhibits a fragmentation pattern consistent with methylation on ring B, corresponding to prunin 4’-methyl ether (commercially known as isosakuranin). For compounds **6**–**10**, a normalized collision energy (NCE) of 35 did not yield effective fragmentation. Therefore, a combination of 30, 40, and 55 NCE was used. For chrysin (**6**, *m/z* 255.0654, [M + H]^+^, RT = 6.09 min), a single monomethylated product (**6a**, RT = 5.66 min) was detected. While MS2 analysis did not provide definitive structural information, its earlier retention time suggests it is likely chrysin 5-methyl ether.^46,51^ When eupatorin (**7**, *m/z* 345.0971, [M + H]^+^, RT = 5.96 min) was used as a substrate, two monomethylated products (**7a** and **7b**) were detected at 5.72 min and 6.21 min. Based on previous reports on HPLC separation, **7a** was identified as eupatorin 5-methyl ether, while **7b** likely corresponds to eupatorin 3’-methyl ether (commercially known as 5-desmethylsinensetin).^52^ For kaempferol (**8**, *m/z* 287.0550, [M + H]^+^, RT = 5.66 min), two monomethylated products (**8a** and **8b**) were detected at 5.37 min and 5.74 min, along with one di-methylated product (**8c**) at 5.32 min. Based on literature, kaempferol 3- and 5-methyl derivatives are prone to CH₄ loss under tandem MS due to their spatial proximity to the C4-keto group, forming a fragment with *m/z* 285 [M + H]^+^. The detection of this fragment in both **8a** and **8b** and their relative retention times suggests that they correspond to kaempferol 5-methyl ether and kaempferol 3-methyl ether, respectively.^50^ The di-methylated product (**8c**) exhibited a distinctive *m/z* 299.0550 fragment, likely resulting from steric and electronic effects that hinder the characteristic Retro-Diels-Alder fragmentation of flavonoids, favoring CH₄ loss instead.^50^ Consequently, we propose it to be kaempferol 3,5-dimethyl ether, further supported by the RT, which is even smaller than that of **8a**. Luteolin (**9**, *m/z* 287.0549, [M + H]^+^, RT = 5.44 min) yielded a monomethylated product (**9a**, RT = 5.67 min), closely matching 5,7-dihydroxy-2-(4-hydroxy-3-methoxyphenyl) chromen-4-one based on the GNPS Library Spectrum CCMSLIB00000079846. Additionally, a di-methylated product (**9b**, RT = 5.29 min) was detected, though the specific methylation sites could not be determined. For amentoflavone (**10**, *m/z* 539.0983, [M + H]^+^, RT = 5.79 min), one monomethylated product (RT = 5.54 min) was detected. MS2 comparison with the GNPS database showed a strong match with amentoflavone 4’-methyl ether (Spectrum ID: CCMSLIB00005724613).

**Figure 7.**
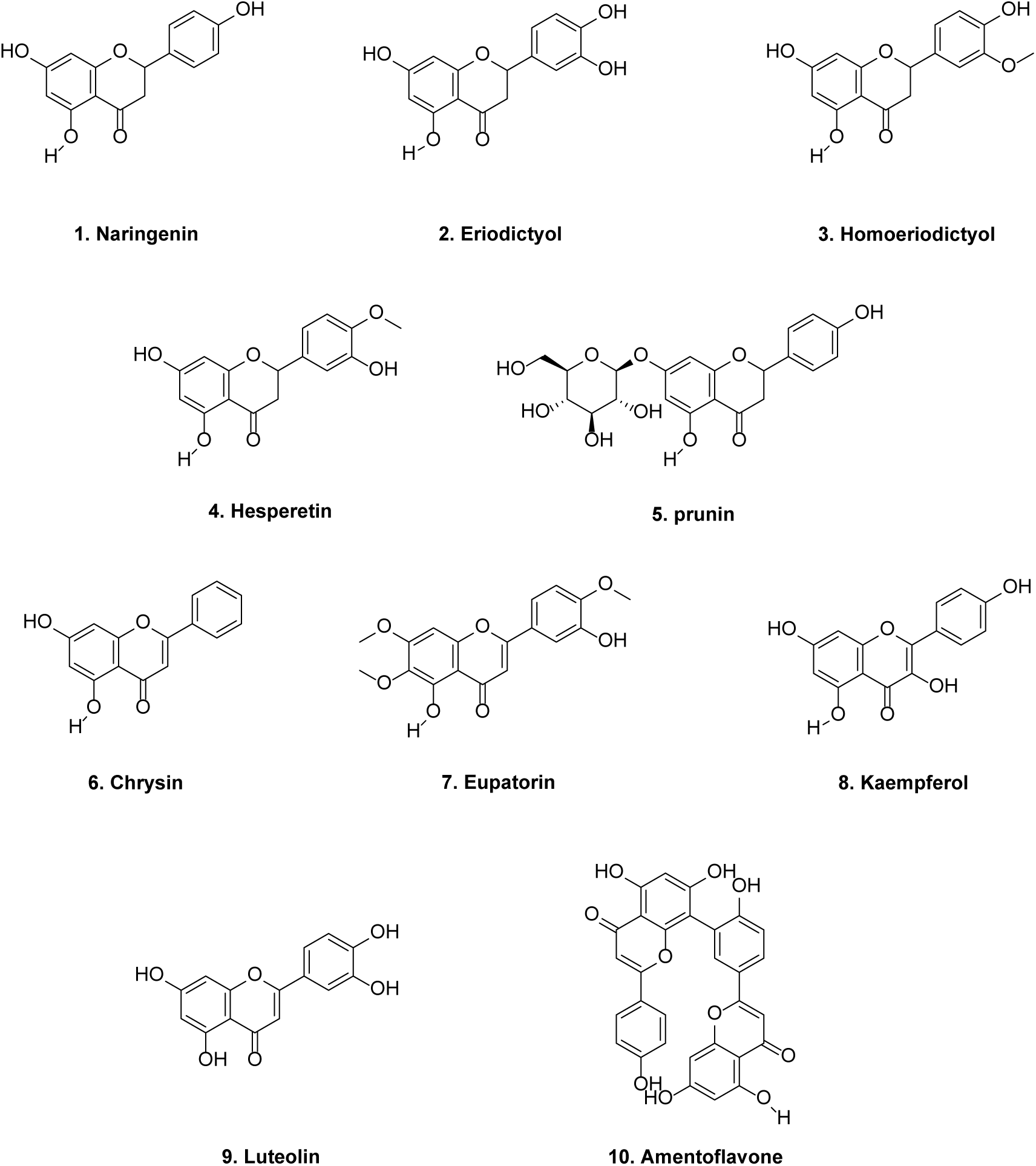
Substrate scope of the methylation reactions catalyzed by StrAOMT wildtype, StrAOMT K212R, and StrAOMT I41L/R171W/K212R. Reaction conditions: 25 mM Tris-HCl (pH 7.5), 2 mM MgCl₂, 1 mM SAM, 500 μM substrate, and 50 μM enzyme, in a final volume of 100 μL, with incubation at 37 °C for 12 hours.

**Table 3.**
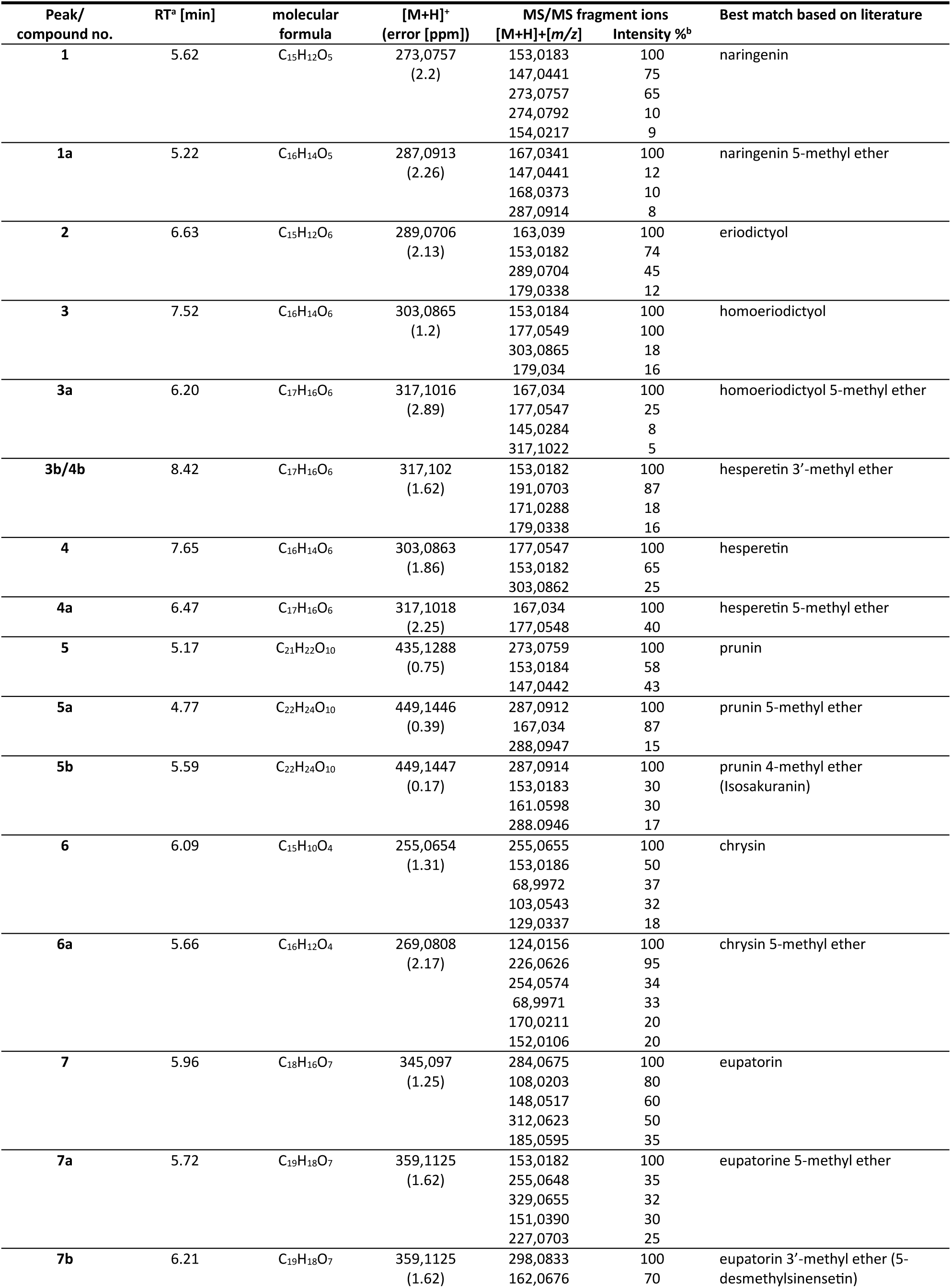
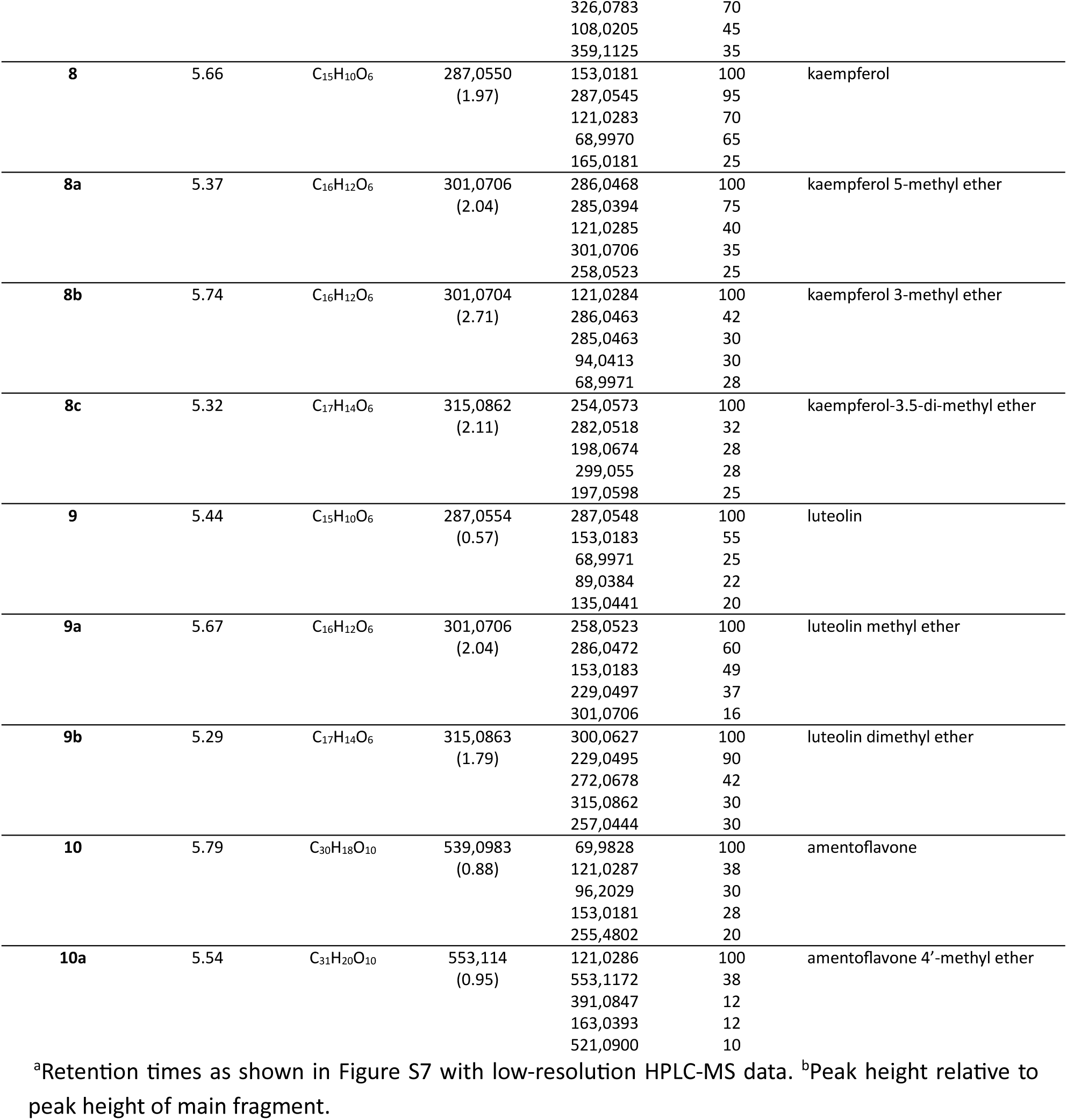
Characteristics of the major peaks detected via HPLC-MS and high-resolution tandem MS in the extracts of enzymatic reactions of the StrAOMT variants.

After putatively identifying the methylated products, we analyzed the product peak areas of the three enzyme variants for a semi-quantitative comparison. This analysis indicates that the triple mutant exhibits enhanced activity towards the formation of naringenin 5-methyl ether, homoeriodictyol 5-methyl ether, hesperetin 5-methyl ether, prunin 5-methyl ether, eupatorin 5-methyl ether, amentoflavone 4’-methyl ether, and luteolin dimethyl ether compared to wild type and double mutant. Notably, it showed a high selectivity for hesperetin 5’-methyl ether formation (>99%). Meanwhile, the double mutant demonstrated increased activity towards chrysin 5-methyl ether and kaempferol dimethyl ether compared to wild type and triple mutant. This suggests that our mutagenesis campaign did influence the regioselectivity of StrAOMT but further engineering is necessary to further enhance the substrate conversion and regioselectivity.

## 4. Discussion

Alpinetin is a naturally occurring 5-O-methylated flavonoid with diverse bioactivities, widely found in medicinal plants such as *Mikania micrantha Kunth*, *Combretum albopunctatum Suess*., *Dalbergia odorifera T.C.Chen*, *Scutellaria indica L*., *Persicaria ferruginea*, and *Alpinia hainanensis K.Schum.*^14,16^ Among these, the *Zingiberaceae* family has the highest levels of alpinetin, which is particularly abundant in its seeds and rhizomes, containing approximately 5 mg per gram of dry biomass. Despite its low abundance in plants, the extraction process relies on solvents such as methanol, ethanol, dichloromethane, and chloroform, which pose environmental and safety concerns. While significant advances have been made in the biosynthesis of pinocembrin, alpinetin biosynthesis remains largely underexplored.^37–41^

In this study, we successfully achieved the *de novo* biosynthesis of alpinetin in *E. coli* for the first time by choice of pathway enzymes, optimizing plasmid configurations, medium composition, and fermentation strategies. Additionally, we enhanced the activity of StrAOMT through directed evolution, resulting in the StrAOMT I41L/R171W/K212R triple mutant. This variant demonstrated an 81-fold increase in alpinetin production *in vitro* compared to the wild-type enzyme. Michaelis-Menten analysis further revealed that the mutant enzyme had a higher turnover number and improved affinity for pinocembrin. Using this improved enzyme, we achieved a final alpinetin titer of approximately 4.98 μM. Interestingly, StrAOMT variants exhibited O-methylation activity toward a broad range of flavonoids, producing valuable 5-O-methylated compounds such as naringenin 5-methyl ether, homoeriodictyol 5-methyl ether, hesperetin 5-methyl ether, prunin 5-methyl ether, chrysin 5-methyl ether, eupatorine 5-methyl ether, and kaempferol 5-methyl ether. Additionally, they produced commercially valuable flavonoids such as isosakuranin and 5-desmethylsinensetin, whose biosynthesis had not been previously reported.^52,53^ Moreover, the StrAOMT I41L/R171W/K212R triple mutant displayed altered regioselectivity toward hesperetin, homoeriodictyol 5-methyl ether, and hesperetin 5-methyl ether formation. These findings highlight the potential of StrAOMT variants for the selective biosynthesis of these valuable O-methylated flavonoids in the future. Recently, the *de novo* biosynthesis of the prenylated flavonoid xanthohumol was reported, with the final titers of xanthohumol and desmethylxanthohumol indicating that the methylation step needs further optimization. Our StrAOMT variants demonstrate significant potential for efficiently methylating this compound.^54^

The final alpinetin titer we achieved is still significantly below the market demand. Swapping the StrAOMT from the plasmid pRSFDuet-1 to pCDFDuet-1 enabled the production of alpinetin, suggesting that there is a pathway burden in the alpinetin biosynthetic pathway. Therefore, improving the pathway flux will be crucial for increasing the final titer of alpinetin. Future efforts should also focus on co-localizing pinocembrin, and StrAOMT variants within the same cellular compartment, for instance by surface display of the OMT^55–57^. Improving the availability of co-substrates for both the chalcone synthase (malonyl-CoA)^58^ and OMT (SAM)^36,59^ should also further enhance the efficiency of the pathway. Although an 81-fold increase in alpinetin production *in vitro* was achieved, the catalytic turnover of the triple mutant is still relatively low and could be further improved by semi-rational engineering. Alternatively, exploring OMTs from *Alpinia* species may offer additional advantages. For alpinetin biosynthesis from phenylalanine, selecting an aromatic amino acid ammonia lyase with low activity toward *p*-coumaric acid but high specificity for phenylalanine would be beneficial. Potential candidates include PpPAL from *Physcomitrella patens subsp. patens*, DdPAL from *Dictyostelium discoideum*, and BlPAL from *Brevibacillus laterosporus*.^42^

In summary, we achieved *de novo* biosynthesis of alpinetin from phenylalanine in *E. coli* for the first time. The engineered StrAOMT variants hold great potential for the selective biosynthesis of valuable O-methylated flavonoids in the future.

## Author contributions

KH and BP conceived the study with contributions from ZW, LZ, and MG; BP cloned plasmids and performed fermentations; BP and KH analyzed biochemical data; BP and LZ performed docking study; KH and BP wrote the manuscript with contributions from ZW, LZ, and MG; all authors have read and approved the final version of the manuscript.

## Conflict of interest

The authors declare no conflict of interest.

## Funding

BP, ZW, and LZ are supported by promotion scholarships from the Chinese Scholarship Council (202008420246, 202309210023, and 202006320070). KH is grateful for funding from the European Union’s Horizon 2020 research and innovation program under the Marie Skłodowska-Curie grant agreement No 893122.

## Acknowledgements

The authors are grateful Dr. Nika Sokolova for advice and support surrounding the OMT work; to Ms Elena Dijkstra and Mr Jorn Pluim, who performed the initial strain characterizations in the framework of their Bachelor research projects; and to the Interfaculty Mass Spectrometry Center of the University of Groningen and the University Medical Center Groningen for their services in high resolution tandem mass spectrometry.

## Notes

### Competing Interest Statement

The authors have declared no competing interest.

## References

(1) Małgorzata Brodowska, K. European Journal of Biological Research Natural Flavonoids: Classification, Potential Role, and Application of Flavonoid Analogues. Eur J Biol Res 2017, 7 (2), 108–123.

(2) Zarebczan, B.; Pinchot, S. N.; Kunnimalaiyaan, M.; Chen, H. Hesperetin, a Potential Therapy for Carcinoid Cancer. Am J Surg 2011, 201 (3), 329–333. 10.1016/j.amjsurg.2010.08.018.

(3) Walle, T.; Ta, N.; Kawamori, T.; Wen, X.; Tsuji, P. A.; Walle, U. K. Cancer Chemopreventive Properties of Orally Bioavailable Flavonoids-Methylated versus Unmethylated Flavones. Biochem Pharmacol 2007, 73 (9), 1288–1296. 10.1016/j.bcp.2006.12.028.

(4) Harmon, A. W.; Patel, Y. M. Naringenin Inhibits Glucose Uptake in MCF-7 Breast Cancer Cells: A Mechanism for Impaired Cellular Proliferation. Breast Cancer Res Treat 2004, 85 (2), 103–110. 10.1023/B:BREA.0000025397.56192.e2.

(5) Gao, Y.; Wang, S. X.; He, L.; Wang, C. L.; Yang, L. Alpinetin Protects Chondrocytes and Exhibits Anti-Inflammatory Effects via the NF-Kappa B/ERK Pathway for Alleviating Osteoarthritis. Inflammation 2020, 43 (5), 1742–1750. 10.1007/s10753-020-01248-3.

(6) Kwon, J. Y.; Jeon, M. T.; Jung, U. J.; Kim, D. W.; Moon, G. J.; Kim, S. R. Perspective: Therapeutic Potential of Flavonoids as Alternative Medicines in Epilepsy. Advances in Nutrition 2019, 10 (5), 778–790. 10.1093/advances/nmz047.

(7) Panche, A. N.; Diwan, A. D.; Chandra, S. R. Flavonoids: An Overview. J Nutr Sci 2016, 5 e47. 10.1017/jns.2016.41.

(8) Zhou, Y.; Ding, Y. L.; Zhang, J. L.; Zhang, P.; Wang, J. Q.; Li, Z. H. Alpinetin Improved High Fat Diet-Induced Non-Alcoholic Fatty Liver Disease (NAFLD) through Improving Oxidative Stress, Inflammatory Response and Lipid Metabolism. BIOMEDICINE & PHARMACOTHERAPY 2018, 97, 1397–1408. 10.1016/j.biopha.2017.10.035.

(9) Hou, S. S.; Yuan, Q.; Cheng, C. R.; Zhang, Z. G.; Guo, B. R.; Yuan, X. X. Alpinetin Delays High-Fat Diet-Aggravated Lung Carcinogenesis. Basic Clin Pharmacol Toxicol 2021, 128 (3), 410. 10.1111/bcpt.13540.

(10) Liu, T. gang; Sha, K. hui; Zhang, L. guo; Liu, X. xian; Yang, F.; Cheng, J. ying. Protective Effects of Alpinetin on Lipopolysaccharide/D-Galactosamine-Induced Liver Injury through Inhibiting Inflammatory and Oxidative Responses. Microb Pathog 2019, 126 (November 2018), 239–244. 10.1016/j.micpath.2018.11.007.

(11) Tan, Y.; Zheng, C. Q. Effects of Alpinetin on Intestinal Barrier Function, Inflammation and Oxidative Stress in Dextran Sulfate Sodium-Induced Ulcerative Colitis Mice. AMERICAN JOURNAL OF THE MEDICAL SCIENCES 2018, 355 (4), 377–386. 10.1016/j.amjms.2018.01.002.

(12) Zeng, C. Y.; Wang, S.; Chen, F. L.; Wang, Z. M.; Li, J. T.; Xie, Z. Y.; Ma, M. J.; Wang, P.; Shen, H. Y.; Wu, Y. F. Alpinetin Alleviates Osteoporosis by Promoting Osteogenic Differentiation in BMSCs by Triggering Autophagy via PKA/MTOR/ULK1 Signaling. PHYTOTHERAPY RESEARCH 2023, 37 (1), 252–270. 10.1002/ptr.7610.

(13) Wikan, N.; Potikanond, S.; Hankittichai, P.; Thaklaewphan, P.; Monkaew, S.; Smith, D. R.; Nimlamool, W. Alpinetin Suppresses Zika Virus-Induced Interleukin-1 Beta Production and Secretion in Human Macrophages. Pharmaceutics 2022, 14 (12), 2800. 10.3390/pharmaceutics14122800.

(14) Gul, S.; Maqbool, M. F.; Zheng, D. Y.; Li, Y. M.; Khan, M.; Ma, T. H. Alpinetin: A Dietary Flavonoid with Diverse Anticancer Effects. Appl Biochem Biotechnol 2022, 194, 4220–4243. 10.1007/s12010-022-03960-2.

(15) Luo, G.; Lin, J.; Cheng, W.; Liu, Z.; Yu, T.; Yang, B. UHPLC-Q-Orbitrap-MS-Based Metabolomics Reveals Chemical Variations of Two Types of Rhizomes of *Polygonatum Sibiricum*. Molecules 2022, 27 (15), 4685. 10.3390/molecules27154685.

(16) Zhao, G.; Tong, Y.; Luan, F.; Zhu, W. J.; Zhan, C. L.; Qin, T. T.; An, W. X.; Zeng, N. Alpinetin: A Review of Its Pharmacology and Pharmacokinetics. Front Pharmacol 2022, 13, 814370. 10.3389/fphar.2022.814370.

(17) Giang, P. M.; Son, P. T.; Matsunami, K.; Otsuka, H. New Diarylheptanoids from *Alpinia Pinnanensis*. Chem Pharm Bull (Tokyo) 2005, 53 (10), 1335–1337. 10.1248/cpb.53.1335.

(18) Bi, S. Y.; Sun, X. Y.; Wang, Y.; Wu, J.; Zhou, H. F. A Sensitive Resonance Rayleigh Light Scattering Method for Alpinetin Using Gold Nanorods Probes. LUMINESCENCE 2018, 33 (7), 1164–1170. 10.1002/bio.3531.

(19) Sokolova, N.; Zhang, L.; Deravi, S.; Oerlemans, R.; Groves, M. R.; Haslinger, K. Structural Characterization and Extended Substrate Scope Analysis of Two Mg2+-Dependent O-Methyltransferases from Bacteria. ChemBioChem 2023, 24 (9), e202300076. 10.1002/cbic.202300076.

(20) Bennett, M. R.; Shepherd, S. A.; Cronin, V. A.; Micklefield, J. Recent Advances in Methyltransferase Biocatalysis. Curr Opin Chem Biol 2017, 37, 97–106. 10.1016/j.cbpa.2017.01.020.

(21) Haslinger, K.; Hackl, T.; Prather, K. L. J. Rapid *in Vitro* Prototyping of O-Methyltransferases for Pathway Applications in *Escherichia Coli*. Cell Chem Biol 2021, 28 (6), 876–886.e4. 10.1016/j.chembiol.2021.04.010.

(22) Peng, B.; Zhang, L.; He, S.; Oerlemans, R.; Quax, W. J.; Groves, M. R.; Haslinger, K. Engineering a Plant Polyketide Synthase for the Biosynthesis of Methylated Flavonoids. J Agric Food Chem 2024, 72 (1), 529–539. 10.1021/acs.jafc.3c06785.

(23) Miyata, S.; Inoue, J.; Shimizu, M.; Sato, R. Xanthohumol Improves Diet-Induced Obesity and Fatty Liver by Suppressing Sterol Regulatory Element-Binding Protein (SREBP) Activation. Journal of Biological Chemistry 2015, 290 (33), 20565–20579. 10.1074/jbc.M115.656975.

(24) Stevens, J. F.; Page, J. E. Xanthohumol and Related Prenylflavonoids from Hops and Beer: To Your Good Health! Phytochemistry 2004, 65 (10), 1317–1330. 10.1016/j.phytochem.2004.04.025.

(25) Struck, A.-W.; Thompson, M. L.; Wong, L. S.; Micklefield, J. S - Adenosyl-Methionine-Dependent Methyltransferases: Highly Versatile Enzymes in Biocatalysis, Biosynthesis and Other Biotechnological Applications. ChemBioChem 2012, 13 (18), 2642–2655. 10.1002/cbic.201200556.

(26) Peng, J.; Liao, C.; Bauer, C.; Seebeck, F. P. Fluorinated S-Adenosylmethionine as a Reagent for Enzyme-Catalyzed Fluoromethylation. Angewandte Chemie - International Edition 2021, 60 (52), 27178–27183. 10.1002/anie.202108802.

(27) Ibrahim, R. K.; Bruneau, A.; Bantignies, B. Plant O-Methyltransferases: Molecular Analysis, Common Signature and Classification. Plant Mol Biol 1998, 36 (1), 1–10. 10.1023/A:1005939803300.

(28) Liu, Y.; Fernie, A. R.; Tohge, T. Diversification of Chemical Structures of Methoxylated Flavonoids and Genes Encoding Flavonoid-O-Methyltransferases. Plants 2022, 11 (4), 564. 10.3390/plants11040564.

(29) Kim, B. G.; Sung, S. H.; Chong, Y.; Lim, Y.; Ahn, J. H. Plant Flavonoid O-Methyltransferases: Substrate Specificity and Application. Journal of Plant Biology 2010, 53 (5), 321–329. 10.1007/s12374-010-9126-7.

(30) Green, M. R.; Sambrook, J. Transformation of *Escherichia Coli* by Electroporation. Cold Spring Harb Protoc 2020, 2020(6) (6), pdb. prot101220. 10.1101/pdb.prot101220.

(31) Larsson, A. AliView: A Fast and Lightweight Alignment Viewer and Editor for Large Datasets. Bioinformatics 2014, 30 (22), 3276–3278. 10.1093/bioinformatics/btu531.

(32) Muhammed, M. T.; Aki-Yalcin, E. Molecular Docking: Principles, Advances, and Its Applications in Drug Discovery. Lett Drug Des Discov 2022, 21 (3), 480–495. 10.2174/1570180819666220922103109.

(33) Çlnaroǧlu, S. S.; Timuçin, E. Comparative Assessment of Seven Docking Programs on a Nonredundant Metalloprotein Subset of the PDBbind Refined. J Chem Inf Model 2019, 59 (9), 3846–3859. 10.1021/acs.jcim.9b00346.

(34) Korb, O.; Stützle, T.; Exner, T. E. Empirical Scoring Functions for Advanced Protein-Ligand Docking with PLANTS. J Chem Inf Model 2009, 49 (1), 84–96. 10.1021/ci800298z.

(35) O’Boyle, N. M.; Banck, M.; James, C. A.; Morley, C.; Vandermeersch, T.; Hutchison, G. R. Open Babel. J Cheminform 2011, 3 (33), 1–14.

(36) Siegrist, J.; Netzer, J.; Mordhorst, S.; Karst, L.; Gerhardt, S.; Einsle, O.; Richter, M.; Andexer, J. N. Functional and Structural Characterisation of a Bacterial O-Methyltransferase and Factors Determining Regioselectivity. FEBS Lett 2017, 591 (2), 312–321. 10.1002/1873-3468.12530.

(37) Tous Mohedano, M.; Mao, J.; Chen, Y. Optimization of Pinocembrin Biosynthesis in *Saccharomyces Cerevisiae*. ACS Synth Biol 2023, 12 (1), 144–152. 10.1021/acssynbio.2c00425.

(38) Kim, B. G.; Lee, H.; Ahn, J. H. Biosynthesis of Pinocembrin from Glucose Using Engineered *Escherichia Coli*. J Microbiol Biotechnol 2014, 24 (11), 1536–1541. 10.4014/jmb.1406.06011.

(39) Dunstan, M. S.; Robinson, C. J.; Jervis, A. J.; Yan, C.; Carbonell, P.; Hollywood, K. A.; Currin, A.; Swainston, N.; Feuvre, R. Le; Micklefield, J.; Faulon, J.-L.; Breitling, R.; Turner, N.; Takano, E.; Scrutton, N. S. Engineering *Escherichia Coli* towards *de Novo* Production of Gatekeeper (2S)-Flavanones: Naringenin, Pinocembrin, Eriodictyol and Homoeriodictyol. Synth Biol 2020, 5 (1), 1–11. 10.1093/synbio/ysaa012.

(40) Wu, J.; Du, G.; Zhou, J.; Chen, J. Metabolic Engineering of *Escherichia Coli* for (2S)-Pinocembrin Production from Glucose by a Modular Metabolic Strategy. Metab Eng 2013, 16 (1), 48–55. 10.1016/j.ymben.2012.11.009.

(41) Wu, J.; Zhang, X.; Zhou, J.; Dong, M. Efficient Biosynthesis of (2S)-Pinocembrin from D-Glucose by Integrating Engineering Central Metabolic Pathways with a PH-Shift Control Strategy. Bioresour Technol 2016, 218, 999–1007. 10.1016/j.biortech.2016.07.066.

(42) Jendresen, C. B.; Stahlhut, S. G.; Li, M.; Gaspar, P.; Siedler, S.; Förster, J.; Maury, J.; Borodina, I.; Nielsen, A. T. Highly Active and Specific Tyrosine Ammonia-Lyases from Diverse Origins Enable Enhanced Production of Aromatic Compounds in Bacteria and *Saccharomyces Cerevisiae*. Appl Environ Microbiol 2015, 81 (13), 4458–4476. 10.1128/AEM.00405-15.

(43) Peng, B.; Dai, L.; Iacovelli, R.; Driessen, A. J. M.; Haslinger, K. Heterologous Naringenin Production in the Filamentous Fungus *Penicillium Rubens*. J Agric Food Chem 2023, 71 (51), 20782–20792. 10.1021/acs.jafc.3c06755.

(44) Chen, H.; Wang, Z.; Cai, H.; Zhou, C. Progress in the Microbial Production of S-Adenosyl-l-Methionine. World J Microbiol Biotechnol 2016, 32 (9), 1–8. 10.1007/s11274-016-2102-8.

(45) Jones, K. L.; Kim, S.-W.; Keasling, J. D. Low-Copy Plasmids Can Perform as Well as or Better Than High-Copy Plasmids for Metabolic Engineering of Bacteria. Metab Eng 2000, 2, 328–338.

(46) Brandt, W.; Manke, K.; Vogt, T. A Catalytic Triad - Lys-Asn-Asp - Is Essential for the Catalysis of the Methyl Transfer in Plant Cation-Dependent O-Methyltransferases. Phytochemistry 2015, 113, 130–139. 10.1016/j.phytochem.2014.12.018.

(47) Kopycki, J. G.; Stubbs, M. T.; Brandt, W.; Hagemann, M.; Porzel, A.; Schmidt, J.; Schliemann, W.; Zenk, M. H.; Vogt, T. Functional and Structural Characterization of a Cation-Dependent O-Methyltransferase from the Cyanobacterium *Synechocystis* Sp. Strain PCC 6803. Journal of Biological Chemistry 2008, 283 (30), 20888–20896. 10.1074/jbc.M801943200.

(48) Patra, N.; Ioannidis, E. I.; Kulik, H. J. Computational Investigation of the Interplay of Substrate Positioning and Reactivity in Catechol O-Methyltransferase. PLoS One 2016, 11 (8), e0161868. 10.1371/journal.pone.0161868.

(49) Adhikari, D.; Gong, D. S.; Oh, S. H.; Sung, E. H.; Lee, S. O.; Kim, D. W.; Oak, M. H.; Kim, H. J. Vasorelaxant Effect of *Boesenbergia Rotunda* and Its Active Ingredients on an Isolated Coronary Artery. Plants 2020, 9 (12), 1–13. 10.3390/plants9121688.

(50) Ma, C.; Lv, H.; Zhang, X.; Chen, Z.; Shi, J.; Lu, M.; Lin, Z. Identification of Regioisomers of Methylated Kaempferol and Quercetin by Ultra High Performance Liquid Chromatography Quadrupole Time-of-Flight (UHPLC-QTOF) Tandem Mass Spectrometry Combined with Diagnostic Fragmentation Pattern Analysis. Anal Chim Acta 2013, 795, 15–24. 10.1016/j.aca.2013.07.038.

(51) Chandran, K. S.; Humphries, J.; Goodger, J. Q. D.; Woodrow, I. E. Molecular Characterisation of Flavanone O-Methylation in *Eucalyptus*. Int J Mol Sci 2022, 23 (6), 3190. 10.3390/ijms23063190.

(52) Liu, R.; Choi, H. S.; Ko, Y. C.; Yun, B. S.; Lee, D. S. 5-Desmethylsinensetin Isolated from Artemisia Princeps Suppresses the Stemness of Breast Cancer Cells via Stat3/IL-6 and Stat3/YAP1 Signaling. Life Sci 2021, 280, 119729. 10.1016/j.lfs.2021.119729.

(53) Sabrin, M. S.; Selenge, E.; Takeda, Y.; Batkhuu, J.; Ogawa, H.; Jamsransuren, D.; Suganuma, K.; Murata, T. Isolation and Evaluation of Virucidal Activities of Flavanone Glycosides and Rosmarinic Acid Derivatives from *Dracocephalum Spp*. against Feline Calicivirus. Phytochemistry 2021, 191, 112896. 10.1016/j.phytochem.2021.112896.

(54) Yang, S.; Chen, R.; Cao, X.; Wang, G.; Zhou, Y. J. *De Novo* Biosynthesis of the Hops Bioactive Flavonoid Xanthohumol in Yeast. Nat Commun 2024, 15 (1), 253. 10.1038/s41467-023-44654-5.

(55) Schüürmann, J.; Quehl, P.; Festel, G.; Jose, J. Bacterial Whole-Cell Biocatalysts by Surface Display of Enzymes: Toward Industrial Application. Appl Microbiol Biotechnol 2014, 98 (19), 8031–8046. 10.1007/s00253-014-5897-y.

(56) Chen, Z.; Duan, R.; Xiao, Y.; Wei, Y.; Zhang, H.; Sun, X.; Wang, S.; Cheng, Y.; Wang, X.; Tong, S.; Yao, Y.; Zhu, C.; Yang, H.; Wang, Y.; Wang, Z. Biodegradation of Highly Crystallized Poly(Ethylene Terephthalate) through Cell Surface Codisplay of Bacterial PETase and Hydrophobin. Nat Commun 2022, 13 (1), 7138. 10.1038/s41467-022-34908-z.

(57) Lozančić, M.; Hossain, A. S.; Mrša, V.; Teparić, R. Surface Display—an Alternative to Classic Enzyme Immobilization. Catalysts 2019, 9 (9), 728. 10.3390/catal9090728.

(58) Klass, S. H.; Wesselkamper, M.; Cowan, A. E.; Lee, N.; Lanclos, N.; Cheong, S.; Wang, Z.; Chen, Y.; Gin, J. W.; Petzold, C. J.; Keasling, J. D. Engineering Controllable Alteration of Malonyl-CoA Levels to Enhance Polyketide Production. Nat Chem Biol 2025, 1–12. 10.1038/s41589-025-01911-6.

(59) Kunjapur, A. M.; Hyun, J. C.; Prather, K. L. J. Deregulation of S-Adenosylmethionine Biosynthesis and Regeneration Improves Methylation in the *E. Coli de Novo* Vanillin Biosynthesis Pathway. Microb Cell Fact 2016, 15 (1), 1–17. 10.1186/s12934-016-0459-x.

