## Supplementary Figures and Tables for "*De novo* biosynthesis of alpinetin enhanced by directed evolution of 5-O-methyltransferase"

b. Genomics for Health in Africa (GHA), Africa-Europe Cluster of Research Excellence (CoRE).

Table of contents:

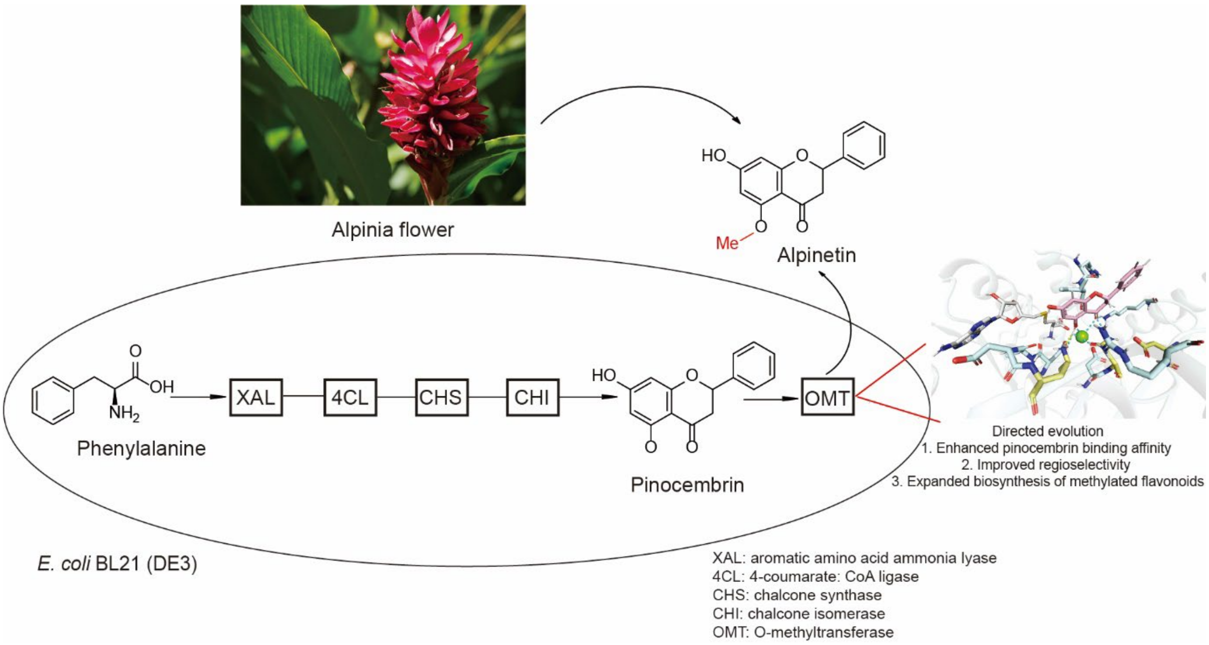

### Supporting materials

#### Sequences of synthetic genes

##### RmXAL (*Rhodotorula mucilaginosa*, GenBank accession number KR095285)

ATGGCACCGAGCGTTGATAGCATTGCAACCAGCGTTGCAAATAGCCTGAGCAATGGTCTGCATGCCGCAGCAGCAGCAAATGGTGGTGAT
GTTCACAAAAAAACCGCAGGCGCAGGTAGCCTGCTGCCGACCACCGAAACCACCCAGCTGGATATTGTTGAACGTATTCTGGCAGATGCC
GGTGCAACCGATCAGATTAAACTGGATGGTTATACCCTGACCCTGGGTGATGTTGTTGGTGCAGCACGTCGTGGTCGTAGCGTTAAAGTTG
CAGATAGTCCGCATATTCGCGAAAAAATTGATGCCAGCGTTGAATTTCTGCGTACCCAACTGGATAATAGCGTTTATGGTGTTACCACCGGTT
TTGGTGGTAGCGCAGATACCCGTACCGAAGATGCAATTAGCCTGCAGAAAGCACTGCTGGAACATCAGCTGTGTGGTGTTCTGCCGACCTC
AATGGATGGTTTTGCACTGGGTCGTGGTCTGGAAAATAGTCTGCCGCTGGAAGTTGTTCGTGGTGCAATGACCATTCGTGTTAATAGTCTG
ACCCGTGGTCATAGTGCAGTTCGTATTGTTGTTCTGGAAGCACTGACCAATTTTCTGAATCATGGTATTACCCCGATTGTTCCGCTGCGTGGC
ACCATTAGCGCAAGCGGTGATCTGAGTCCGCTGAGCTATATTGCAGCAAGCATTACCGGTCATCCGGATAGCAAAGTTCATGTTGATGGCAA
AATTATGAGCGCACAAGAAGCAATTGCACTGAAAGGTCTGCAGCCGGTTGTGCTGGGTCCGAAAGAAGGTCTGGGTCTGGTTAATGGCAC
CGCAGTTAGCGCCAGCATGGCAACCCTGGCACTGACCGATGCACATGTTCTGAGCCTGCTGGCACAGGCCCTGACCGCACTGACAGTTGA
AGCAATGGTTGGTCATGCAGGTAGCTTTCATCCGTTTCTGCATGATGTTACCCGTCCGCATCCGACCCAGATTGAAGTTGCACGTAATATTCG
TACCCTGCTGGAAGGTAGCAAATATGCAGTTCATCATGAAACCGAGGTGAAAGTGAAAGATGATGAAGGTATTCTGCGTCAGGATCGTTAT
CCGCTGCGCTGTAGTCCGCAGTGGCTGGGTCCTCTGGTTAGCGATATGATTCATGCACATGCAGTGCTGAGCCTGGAAGCAGGTCAGAGC
ACCACCGATAATCCGCTGATTGATCTGGAAAACAAAATGACCCATCATGGTGGTGCATTTATGGCAAGCAGCGTTGGCAATACCATGGAAA
AAACCCGTCTGGCAGTTGCACTGATGGGTAAAGTTAGTTTTACCCAGCTGACCGAAATGCTGAATGCAGGTATGAATCGTGCACTGCCGAG
CTGTCTGGCAGCAGAAGATCCGAGCCTGAGTTATCATTGTAAAGGTCTGGATATCGCAGCAGCCGCATATACCAGCGAACTGGGTCATCTG
GCAAATCCGGTTAGCACCCATGTTCAGCCTGCCGAAATGGGTAATCAGGCAATTAATTCACTGGCCCTGATTAGCGCACGTCGTACAGCCG
AAGCAAATGATGTTCTGTCACTGCTGCTGGCAACCCATCTGTATTGTGTGCTGCAGGCAGTTGATCTGCGTGCAATGGAATTTGAACATACC
AAAGCATTTGAACCGATGGTTACAGAACTGCTGAAACAGCATTTTGGTGCACTGGCAACCGCAGAAGTTGAAGATAAAGTTCGTAAAAGC
ATCTACAAACGCCTGCAACAGAACAATAGCTACGATCTGGAACAGCGCTGGCATGATACCTTTAGCGTTGCCACCGGTGCAGTTGTTGAAG
CACTGGCAGGTCAAGAAGTTAGCCTGGCAAGCCTGAATGCATGGAAAGTTGCATGTGCCGAAAAAGCCATTGCCCTGACCCGTAGCGTTC
GTGATAGCTTTTGGGCAGCACCGAGCAGCAGCTCACCGGCACTGAAATATCTGTCACCGCGTACCCGTGTTCTGTATAGCTTTGTTCGTGAA
GAAGTTGGCGTTAAAGCCCGTCGCGGTGATGTTTATCTGGGTAAACAAGAAGTGACCATTGGTACAAATGTGAGCCGTATTTATGAAGCCA
TTAAAAGCGGTTGTATTGCACCGGTTCTGGTTAAAATGATGGCCAAGCTTGCGGCCGCATAA

##### Gm4CL (*Glycine max*, GenBank accession number X69955)

ATGATTACCCTGGCACCGAGTCTTGATACCCCGAAAACCGATCAGAATCAGGTGAGCGATCCGCAGACCAGCCATGTGTTTAAATCGAAAC
TGCCGGATATCCCGATTAGCAACCATCTGCCGCTGCACAGCTACTGCTTCCAGAACCTGAGCCAGTTTGCCCATCGCCCGTGCCTGATTGTG
GGTCCGGCCAGCAAAACCTTCACCTATGCGGATACCCACCTGATTAGCTCAAAAATTGCGGCGGGTCTGAGCAACCTTGGCATCCTGAAAG
GCGATGTGGTGATGATTCTGCTGCAGAATAGCGCGGATTTCGTGTTTTCCTTTCTGGCGATTAGCATGATTGGCGCGGTCGCGACCACCGC
GAATCCGTTTTACACCGCGCCGGAAATTTTTAAACAGTTTACGGTGAGCAAAGCCAAACTGATTATCACCCAGGCGATGTATGTGGATAAAC
TGCGCAATCACGATGGAGCCAAACTGGGCGAAGATTTTAAAGTTGTGACCGTGGATGATCCGCCGGAAAACTGCCTGCATTTTAGCGTTCT
GAGCGAAGCCAACGAAAGCGATGTGCCGGAAGTTGAAATTCATCCGGATGATGCGGTCGCCATGCCGTTCAGCAGCGGTACCACCGGTCT
GCCGAAAGGCGTGATTCTGACCCATAAAAGCCTGACCACGAGCGTGGCCCAGCAGGTGGATGGCGAAAACCCGAACCTGTATCTGACCAC
CGAAGATGTTCTGCTGTGTGTACTGCCGCTGTTTCATATTTTCAGCTTGAACAGCGTGCTGCTGTGTGCGCTGCGCGCGGGCAGTGCGGTG
CTGCTGATGCAGAAATTCGAAATTGGCACCCTGCTGGAACTGATTCAGCGTCATCGTGTCAGCGTTGCAATGGTTGTGCCGCCGCTGGTTC
TGGCGCTGGCCAAAAATCCGATGGTGGCGGATTTTGACCTGAGCAGCATTCGCCTGGTGCTGTCGGGCGCGGCACCGCTGGGCAAAGAA
CTGGAAGAGGCACTGCGTAATCGTATGCCGCAGGCGGTTCTGGGCCAGGGCTACGGCATGACCGAAGCGGGCCCGGTGCTGAGCATGTG
TCTGGGCTTCGCCAAACAGCCGTTTCAGACCAAAAGCGGCAGCTGCGGCACCGTGGTGCGCAATGCCGAACTGAAAGTGGTGGATCCGG
AAACCGGCCGCAGCCTGGGCTATAACCAGCCGGGCGAAATTTGTATTCGCGGCCAGCAAATTATGAAAGGCTATCTGAATGACGAAGCCG
CAACCGCCTCAACCATTGATAGCGAAGGCTGGCTGCATACCGGCGATGTGGGCTATGTTGATGATGATGATGAAATTTTTATTGTGGATCGC
GTGAAAGAACTGATTAAATATAAAGGCTTTCAGGTGCCGCCGGCGGAACTGGAAGGCCTGCTGGTGAGCCATCCGAGCATTGCGGATGCG
GCGGTGGTGCCGCAGAAAGATGTGGCGGCCGGTGAAGTGCCGGTGGCGTTTGTGGTGCGCAGCAACGGTTTTGATCTGACCGAAGAAG
CCGTAAAAGAATTTATTGCAAAACAGGTAGTGTTCTATAAACGCCTGCATAAAGTGTATTTTGTTCATGCCATTCCGAAAAGCCCGAGCGGC
AAAATTCTGCGCAAAGATCTGCGTGCGAAACTGGAAACCGCCGCGACCCAGACCCCGTAA

##### CsCHS (*Camellia sinensis*, GenBank accession number D26593):

ATGGTGACCGTGGAAGATATTCGCCGTGCGCAGCGCGCCGAAGGCCCGGCGACCGTTATGGCGATTGGCACGGCGACCCCGCCGAATTG
CGTGGATCAGAGCACCTACCCGGATTATTACTTTCGCATTACCAACAGCGAACACAAAGCGGAACTGAAAGAAAAATTTAAACGCATGTGC
GATAAAAGCATGATTAAAAAACGTTATATGTATCTGACCGAAGAAATTCTGAAAGAAAACCCGCAGGTGTGTGAATATATGGCCCCGAGTCT
GGATGCCCGCCAGGATATGGTGGTCGTGGAAGTGCCGAAACTGGGCAAAGAAGCCGCGACGAAAGCCATCAAAGAATGGGGTCAGCCG
AAATCTAAAATTACGCACCTGGTTTTTTGTACCACCTCAGGCGTTGATATGCCGGGCGCGGATTACCAGCTGACCAAACTGCTGGGTCTGCG
TCCGTCTGTGAAACGCCTGATGATGTATCAGCAGGGCTGCTTCGCGGGCGGCACCGTGCTGCGTCTGGCGAAAGATCTGGCGGAAAACAA

TAAAGGCGCCCGCGTGCTGGTGGTGTGCAGCGAAATTACCGCGGTGACCTTTCGTGGCCCGAGCGATACCCACCTGGATAGCCTGGTGGG
TCAGGCGCTGTTTGGTGACGGCGCCGCGGCGATTATTGTTGGCAGCGATCCGATTCCGGAAGTGGAAAAACCGCTGTTCGAACTGGTGTC
AGCGGCGCAGACCATTCTGCCGGATAGCGATGGCGCGATCGATGGTCATCTGCGCGAAGTGGGCCTGACCTTTCACCTGCTGAAAGATGT
TCCGGGCCTGATTAGCAAAAACATTGAAAAAAGCCTGGCGGAAGCGTTCCAGCCGCTGGGCATCAGCGATTGGAACAGCCTTTTCTGGAT
TGCGCATCCGGGTGGCCCGGCGATTCTGGATCAGGTGGAACTGAAACTGGGCTTAAAAGAAGAAAAACTGCGCGCGACTCGCCACGTGC
TGAGCGAATATGGCAACATGAGCAGCGCCTGCGTGCTGTTCATTCTGGATGAAATGCGCAAAAAAAGCGCAGCCGATGGCCTGAAAACCA
CGGGCGAAGGCCTGGAATGGGGCGTGCTGTTTGGCTTTGGCCCGGGCCTGACGGTGGAAACCGTGGTGCTGCATAGCCTGAGCACCTAA

### Supporting Tables

**Table S1**. Primers used in this study.

| Primers | Primer sequences |
|---|---|
| BPE_41NNK_FP | ggggccacattmnnatgaggcagaccggctg |
| BPE_41NNK_RP | gtctgcctcatnnkaatgtggccccgaacca |
| BPE_42NNK_FP | ggccacmnntatatgaggcagaccggctgc |
| BPE_42NNK_RP | ggtctgcctcatatannkgtggccccgaac |
| BPE_171NNK_FP | gtaactcccccaccmnngactacattgtctacgacga |
| BPE_171NNK_RP | tagacaatgtagtcnnkggtgggggagttacagatgc |
| BPE_212NNK_FP | aaccgtcgtaaccmnnggaacccaccgtctgaacc |
| BPE_212NNK_RP | gacggtgggttccnnkggttacgacggttttgctt |

### Supporting Figures

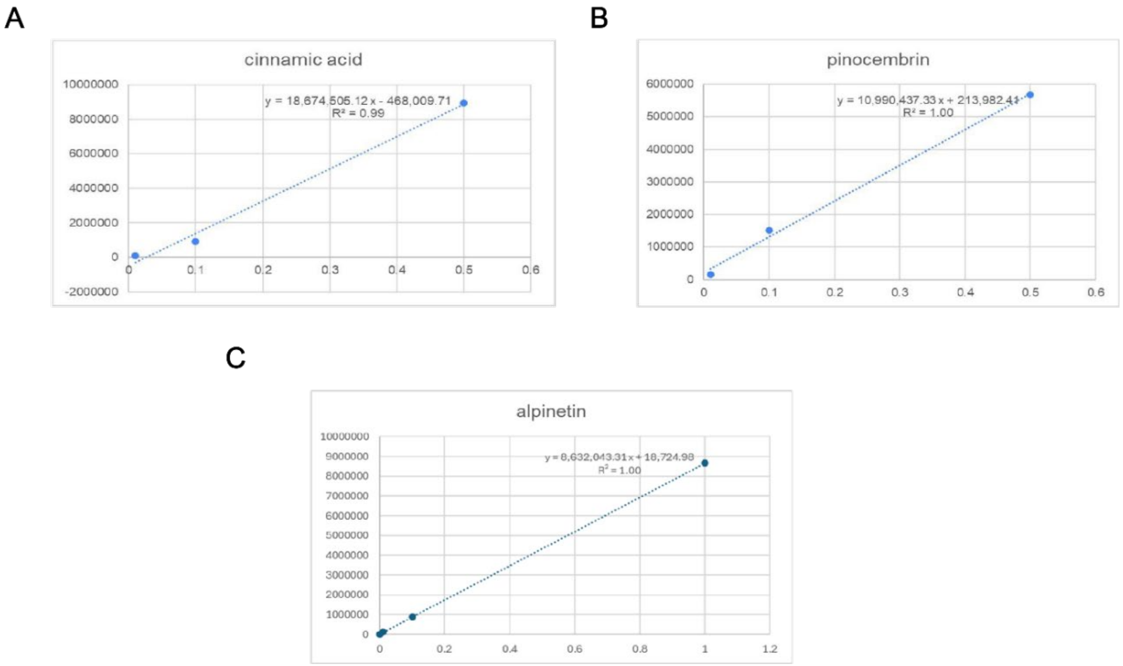


**Figure S1**. Calibration plot of cinnamic acid (A), pinocembrin (B), and alpinetin (C) dissolved in DMSO and analyzed by HPLC. The compounds were detected at 288 nm and the analysis was performed as described in the method section. The range of calibration curve was 0.01 mM to 0.5 mM.

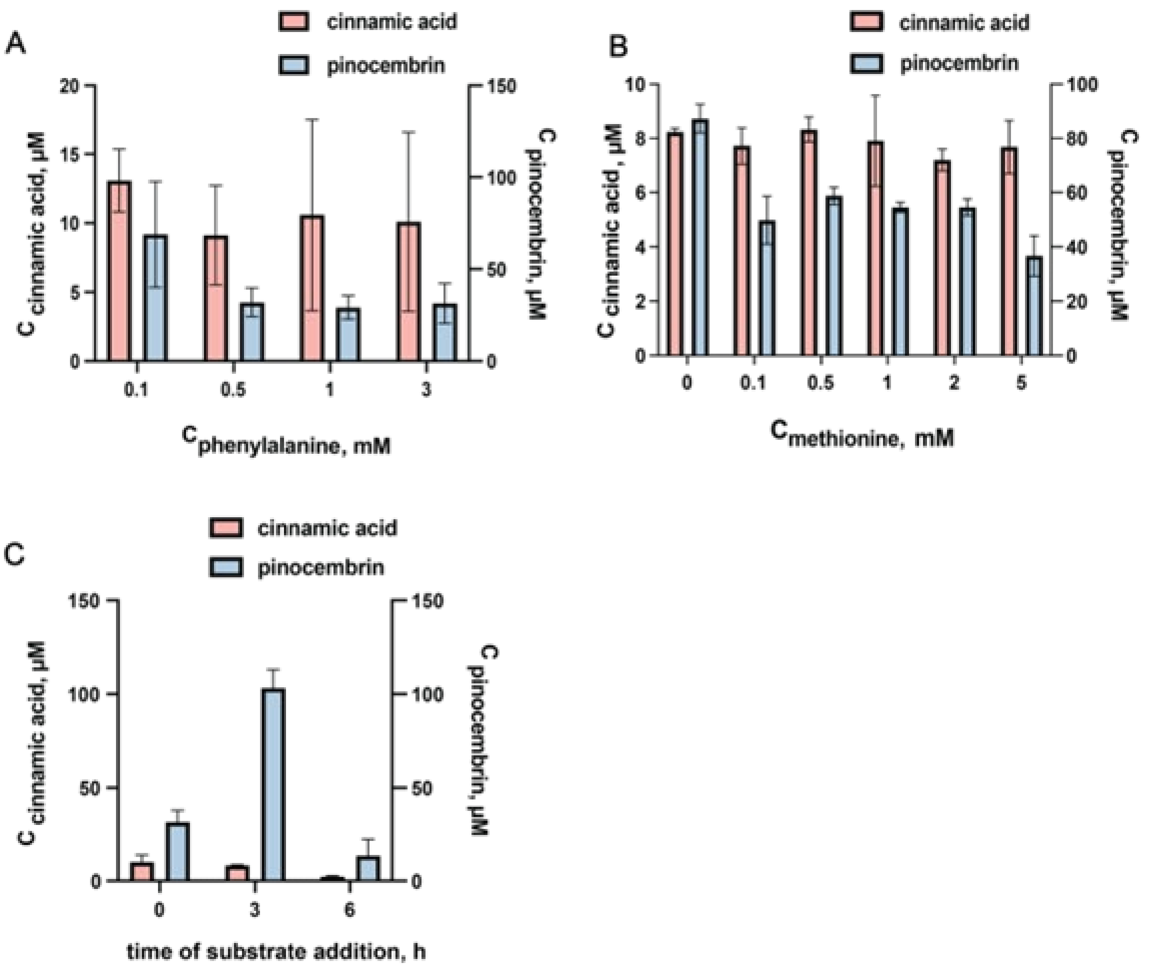


**Figure S2**. Fermentation of *E. coli* MG1655 (DE3) harboring the flavonoid biosynthesis pathway. Titers of cinnamic acid (pink, left axis) and pinocembrin (blue, right axis) were determined 48 h after induction of enzyme expression. A) Cinnamic acid and pinocembrin concentration upon feeding with different concentration of phenylalanine. B) Cinnamic acid and pinocembrin concentration upon feeding with different concentration of methionine. A) Cinnamic acid and pinocembrin concentration with varying substrate addition time. (bars represent mean +/-SD, n=3)

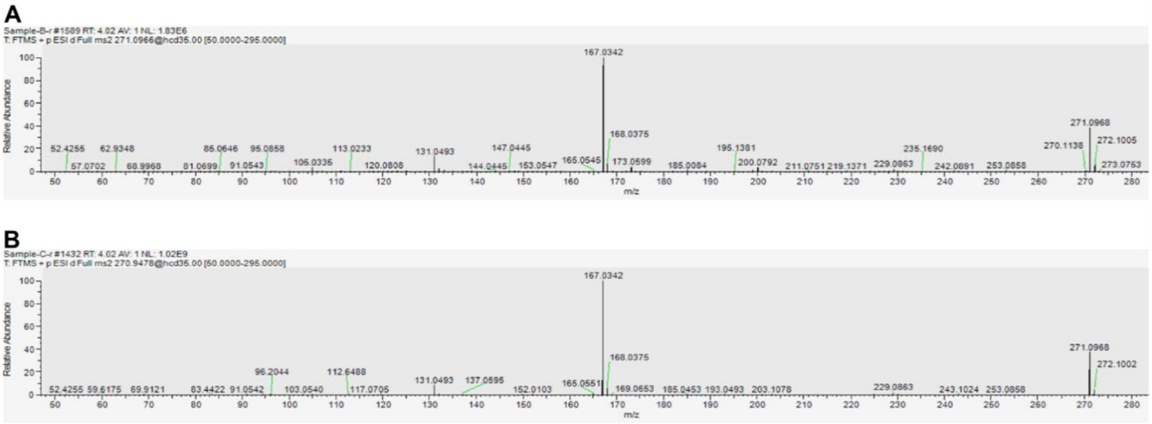


**Figure S3**. Product ion mass spectra (MS2) in high-resolution tandem MS of: A) alpinetin extracted from the S7 fermentation culture (m/z 271.0968, RT=4.02 min), B) commercial standard of alpinetin (m/z 271.0968, RT=4.02 min).

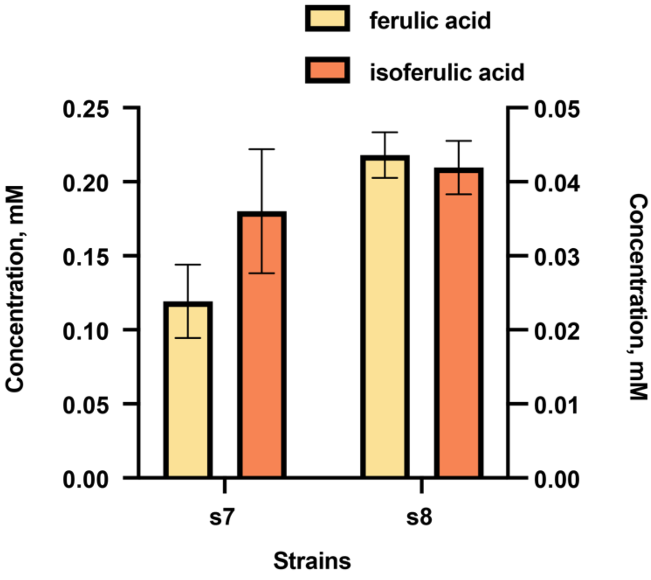


**Figure S4**. Biotransformation of fed caffeic acid (1 mM) in resting *E. coli* BL21 (DE3) strains harboring the flavonoid biosynthesis pathway. Bars represent mean +/- SD, n=3.

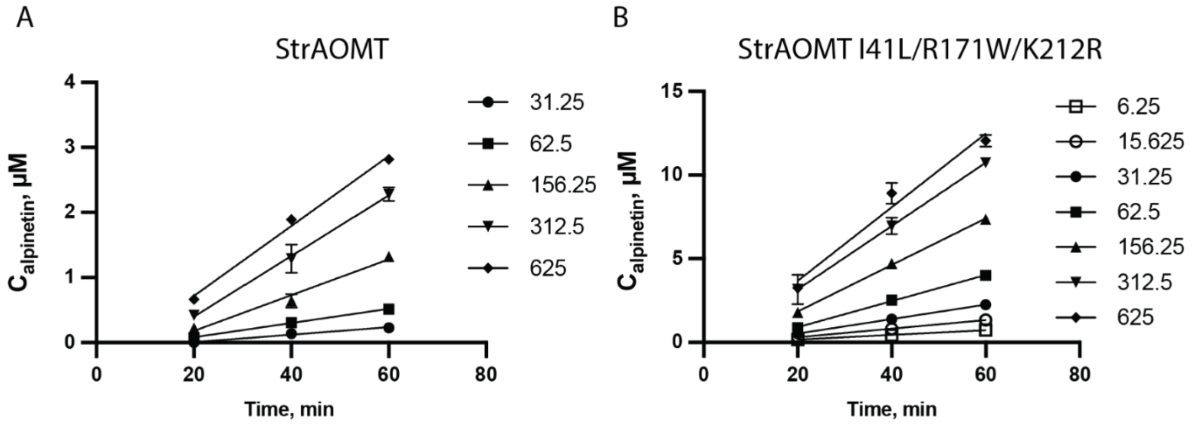


**Figure S5.** Time progress curves underlying the steady-state kinetics analysis. Alpinetin concentration obtained in *in vitro* turnovers catalyzed by A) StrAOMT wild type and B) StrAOMT I41L/R171W/K212R in the presence of varying concentrations of pinocembrin (6.25, 15.625, 31.25, 62.5, 156.25, 312.5, and 625 μM) and a fixed concentration of SAM (1 mM). Data points represent mean +/- SD, n=3, line represents linear regression to determine the apparent initial velocities for each substrate concentration. Replots of initial velocities are shown in Figure 4C and D.

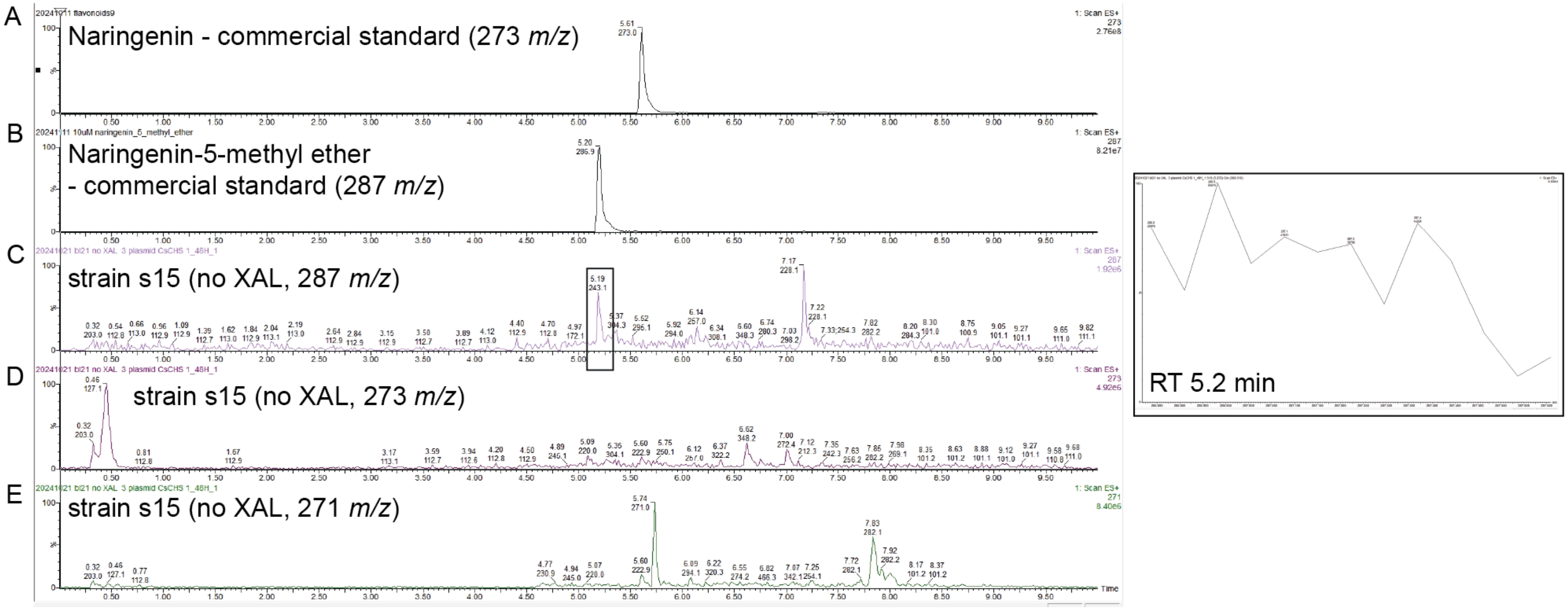


**Figure S6**. Extracted ion chromatograms of: (A) Naringenin standard compound (m/z 273 $[M+H]^+$), (B) Naringenin 5-methyl ether standard compound (m/z 287 $[M+H]^+$), (C) Naringenin 5-methyl ether obtained from s15 fermentation broth, (D) Naringenin obtained from s15 fermentation broth, (E) Alpintin (m/z 271 $[M+H]^+$) detected in s15 fermentation broth, (F) Mass spectrum corresponding to the peak at m/z 243.1 $[M+H]^+$.

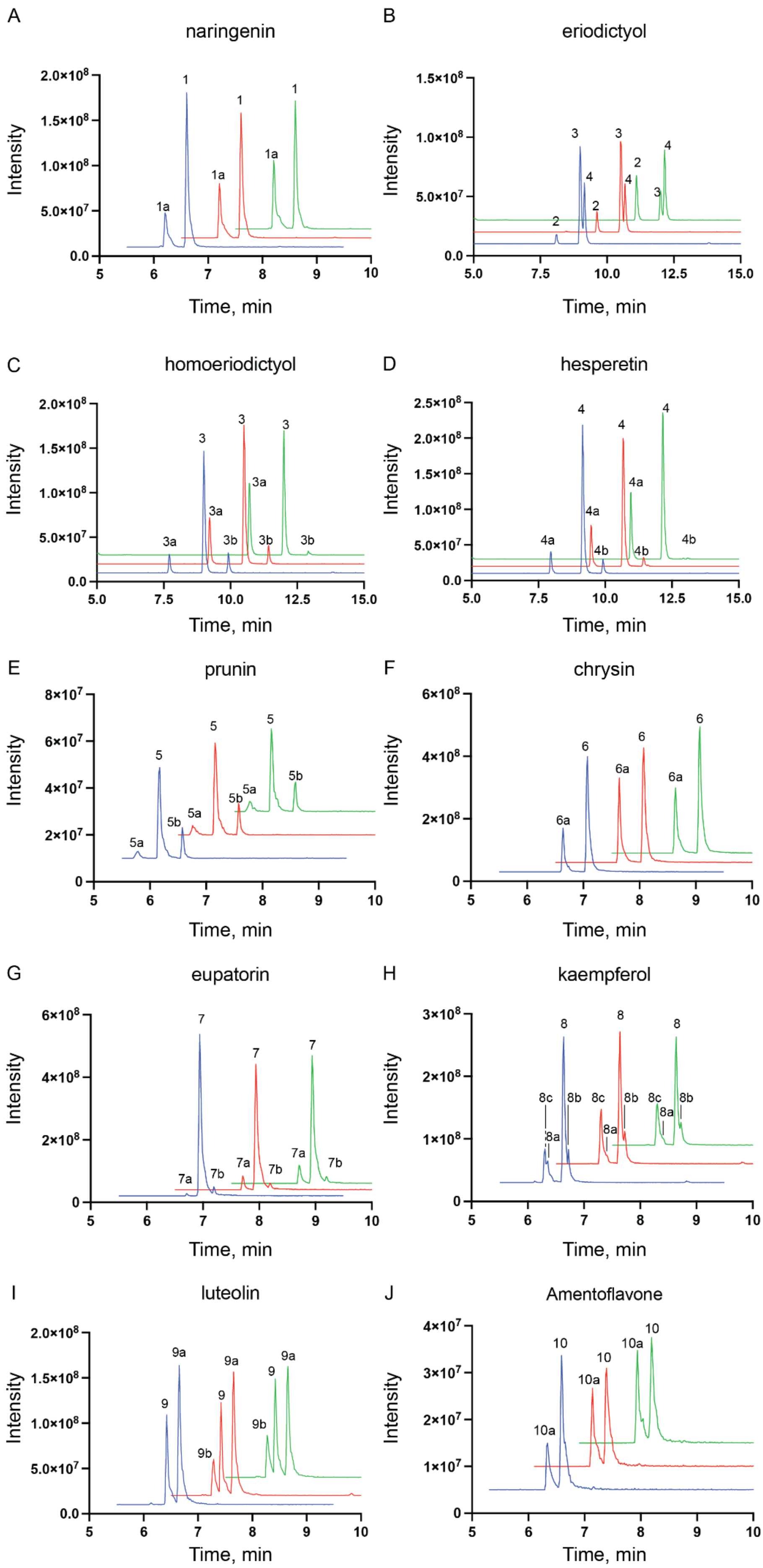
A
naringenin
B
eriodictyol
C
homoeriodictyol
D
hesperetin
E
prunin
F
chrysin
G
eupatorin
H
kaempferol
I
luteolin
J
Amentoflavone
Intensity
Time, min

**Figure S7.** Extracted ion chromatograms of the methylated products from enzymatic reactions by StrAOMT variants analyzed by HPLC-MS. Traces of the three enzyme variants shown with an x/y offset for clarity. Blue traces: StrAOMT wildtype, red traces: StrAOMT K212R, green traces: StrAOMT I41L/R171W/K212R.

1. naringenin

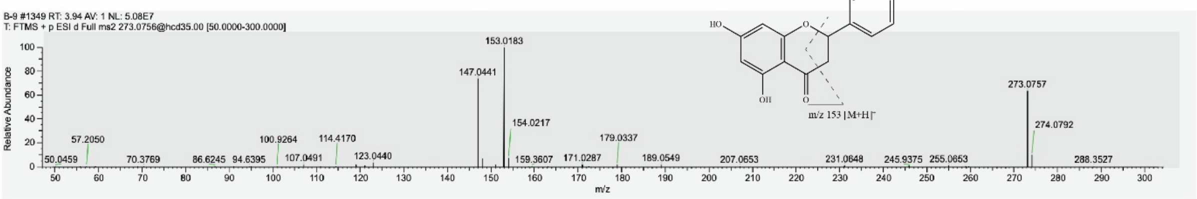


1a. naringenin 5-methyl ether

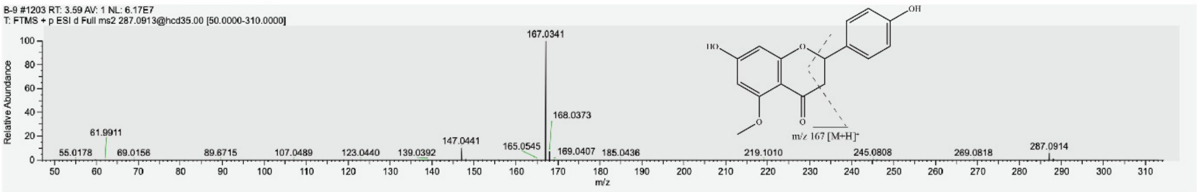


2. eriodictyol

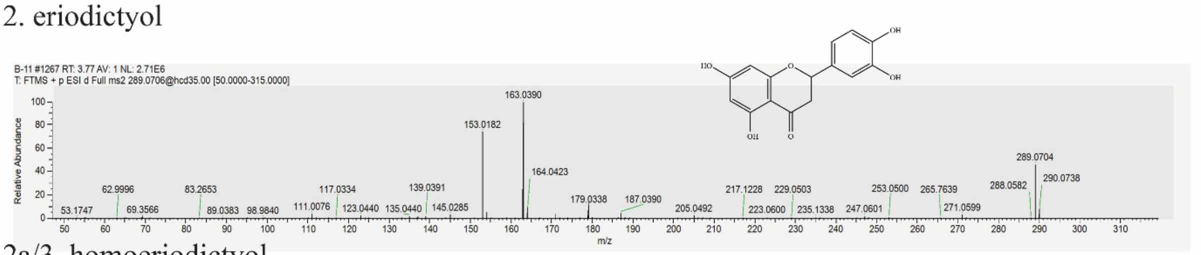


2a/3. homoeriodictyol

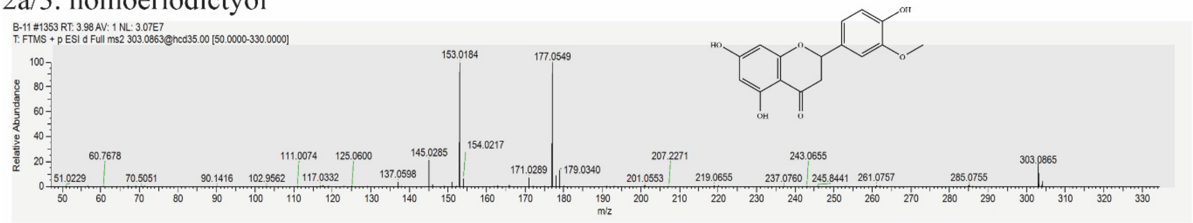


2b/4. hesperetin

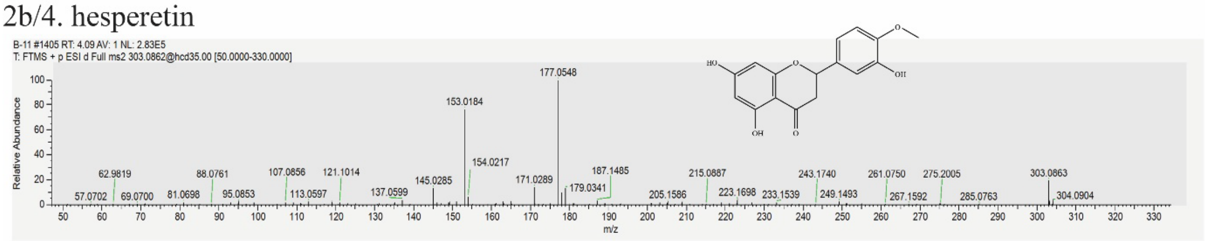

3. homoeriodictyol

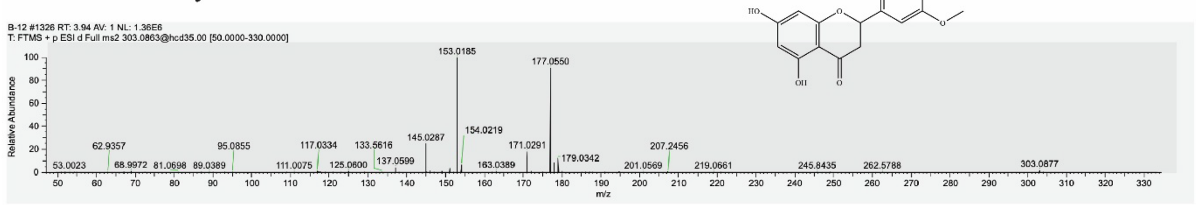


3a. homoeriodictyol 5-methyl ether

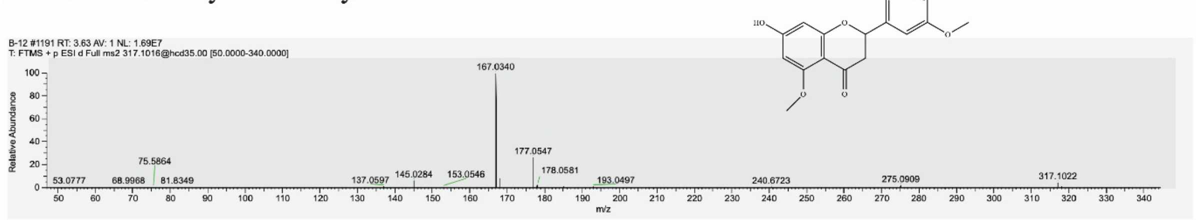


3b. hesperetin 3'-methyl ether

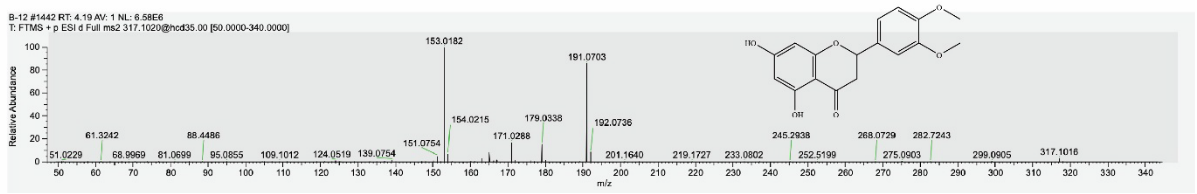


4. hesperetin

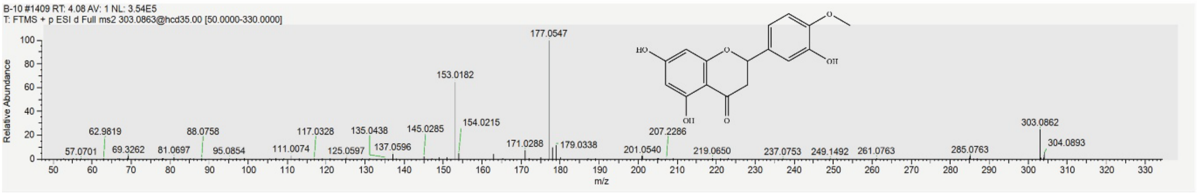


4a. hesperetin 5-methyl ether

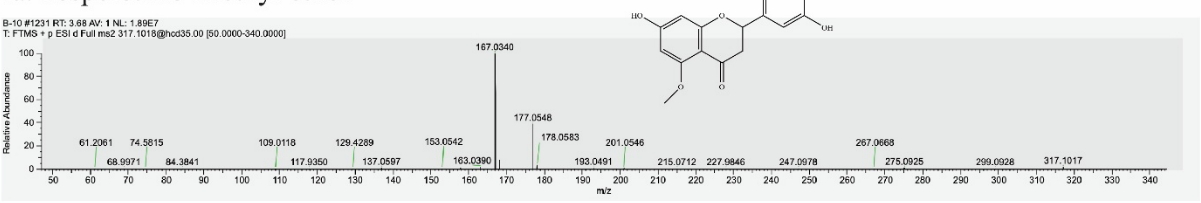


4b. hesperetin 3'-methyl ether

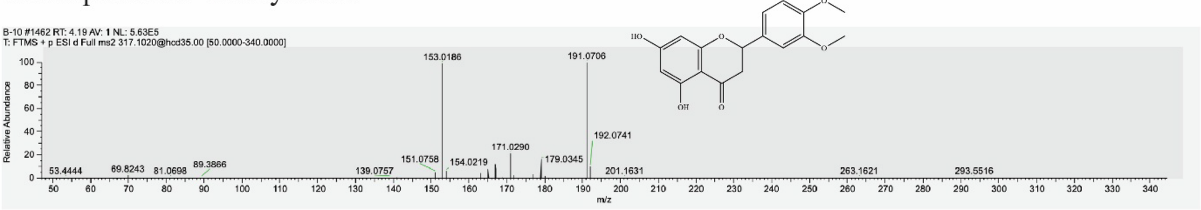

5. prunin

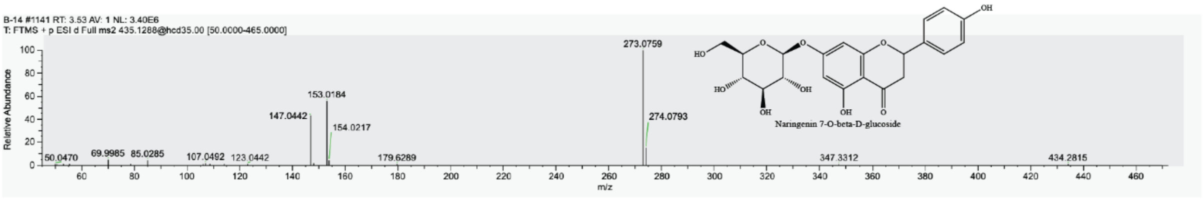


5a. prunin 5-methyl ether

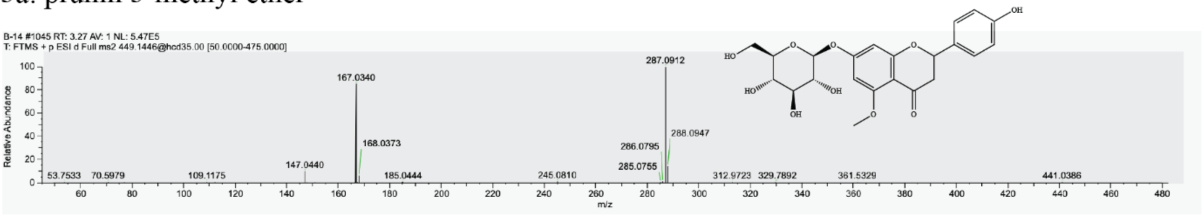


5b. Isosakuranin, prunin 4’-methyl ether

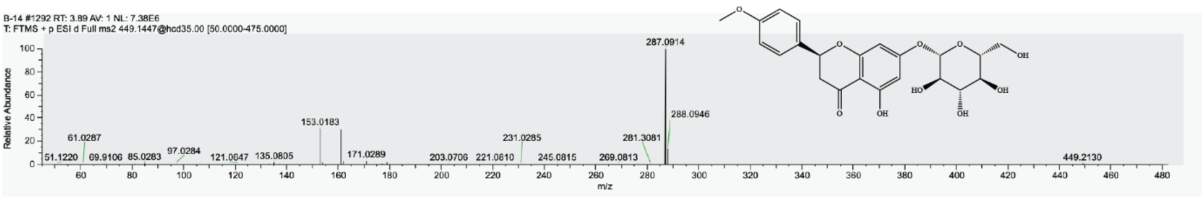


6. chrysin

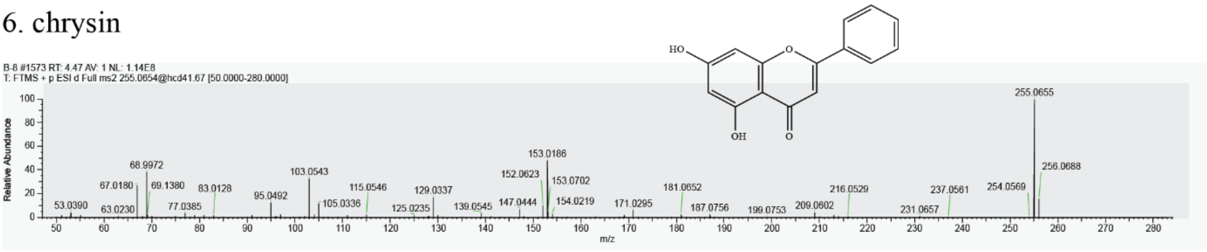


6a. chrysin 5-methyl ether

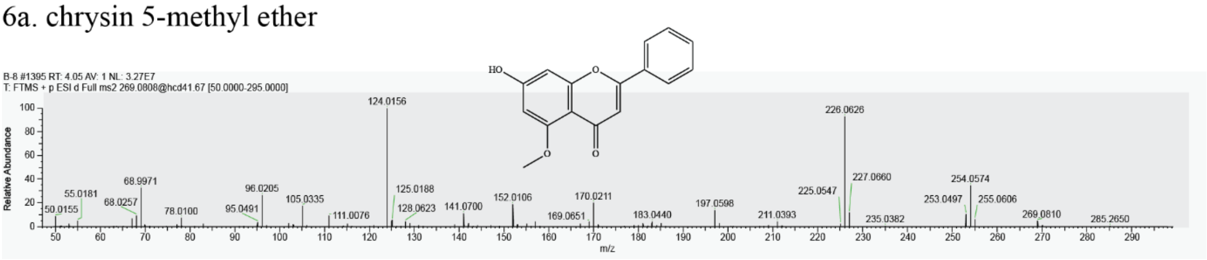

#### 7. eupatorin

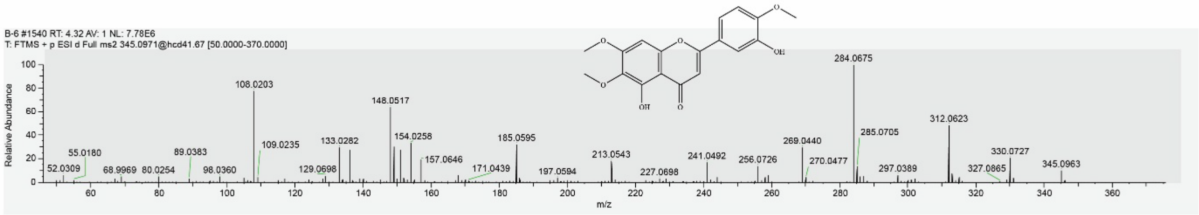


#### 7a. eupatorin 5-methyl ether

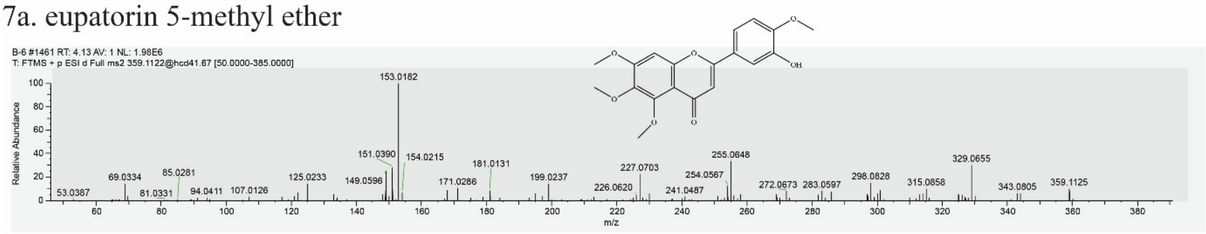


#### 7b. eupatorin 3'-methyl ether, 5-Desmethylsinensetin

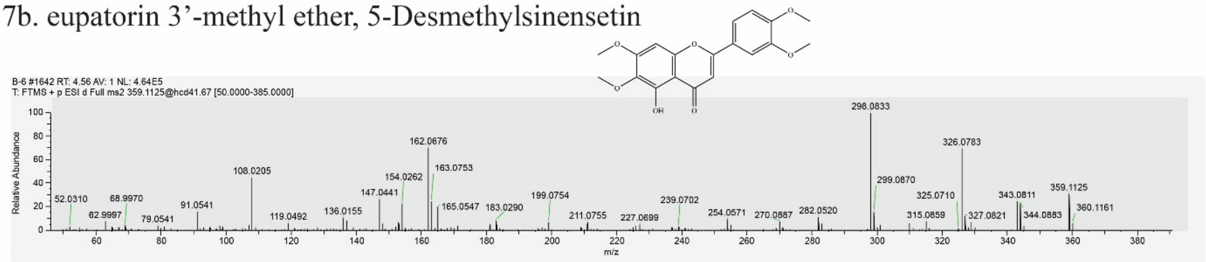


#### 8. kaempferol

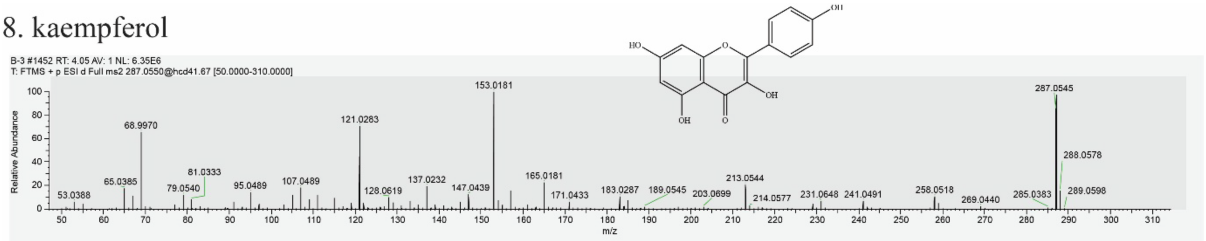


#### 8a. kaempferol 5-methyl ether

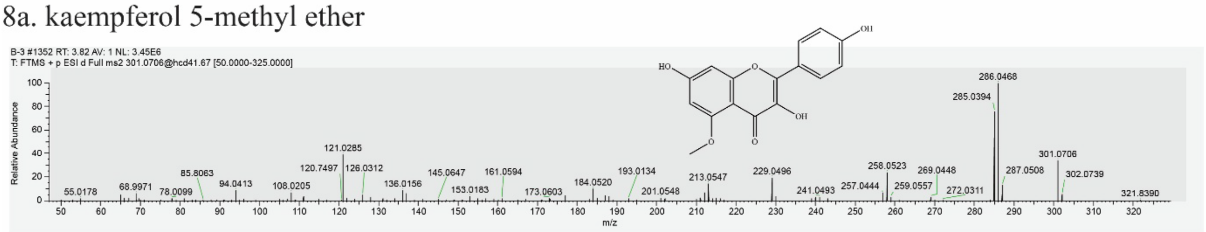


#### 8b. kaempferol 3-methyl ether

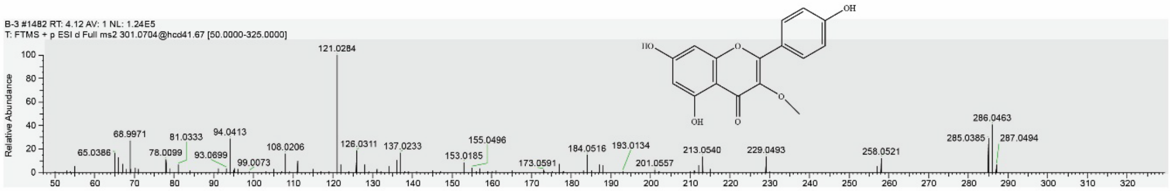


#### 8c. kaempferol 3.5-di-methyl ether

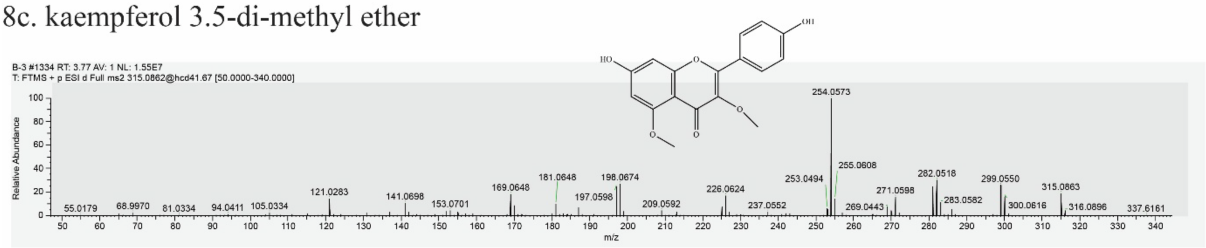

9. luteolin

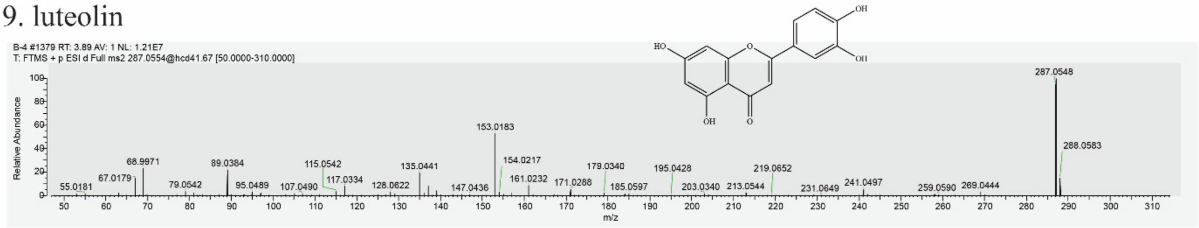


9a. luteolin methyl ether

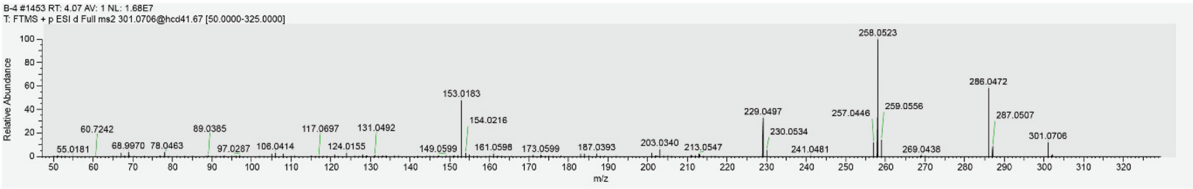


9b. luteolin di-methyl ether

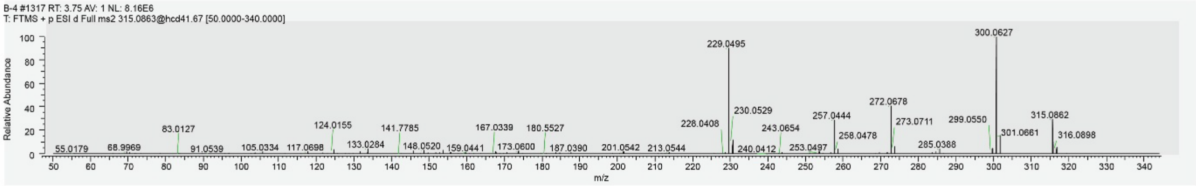


10. amentoflavone

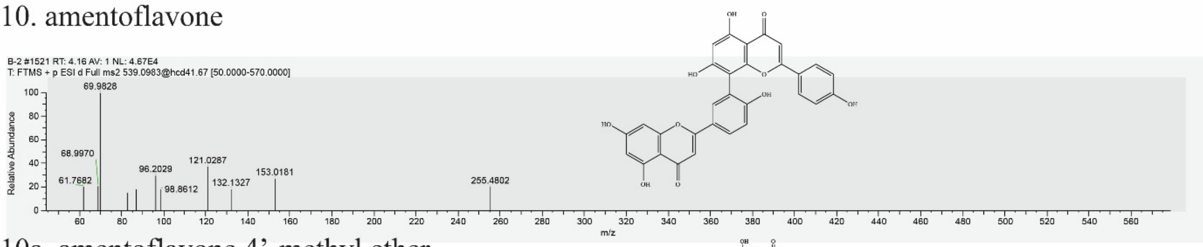


10a. amentoflavone 4’-methyl ether

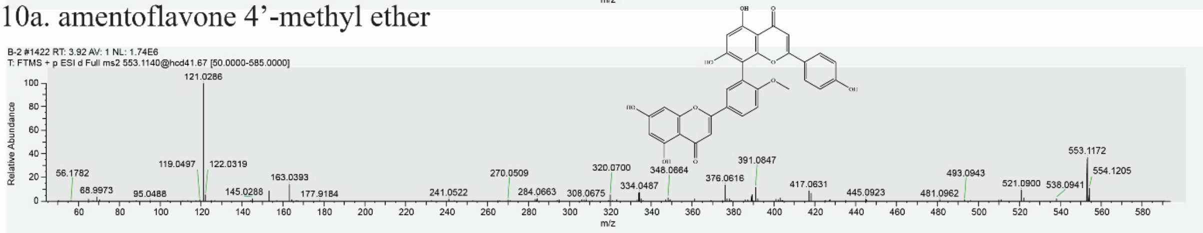


**Figure S8**. Ion mass spectra (MS2) of methylated flavonoids produced by StrAOMT variants, obtained through high-resolution tandem MS analysis.
